# Fillable and Unfillable Gaps in Rice Transcriptome under Field and Controlled Environments

**DOI:** 10.1101/2021.07.31.454577

**Authors:** Yoichi Hashida, Ayumi Tezuka, Yasuyuki Nomura, Mari Kamitani, Makoto Kashima, Yuko Kurita, Atsushi J. Nagano

**Affiliations:** Faculty of Agriculture, Takasaki University of Health and Welfare, Takasaki, Gunma, Japan; Research Institute for Food and Agriculture, Ryukoku University, Otsu, Shiga, Japan; Faculty of Agriculture, Ryukoku University, Otsu, Shiga, Japan; College of Science and Engineering, Aoyama Gakuin University, Sagamihara, Kanagawa, Japan; Institute for Advanced Biosciences, Keio University, Tsuruoka, Yamagata, Japan

**Author notes:** Corresponding author: Atsushi J. Nagano, Postal Address: Faculty of Agriculture, Ryukoku University, Yokotani 1-5, Seta Ohe-cho, Otsu, Shiga 520-2194, Japan.

**Keywords:** Rice, Circadian clock, Field, Growth chamber, Pathogen defence, Phenylpropanoid metabolism, RNA-Seq, Transcriptome, Sugar metabolism

## Abstract

The differences between plants grown in field and controlled environments have long been recognised; however, few studies have addressed the underlying molecular mechanisms. Here, we show fillable and unfillable gaps in the transcriptomes of rice grown in field and controlled environments by utilising SmartGC, a high-performance growth chamber that reproduces the fluctuating irradiance, temperature, and humidity of field environments. Rice transcriptome dynamics in SmartGC mimicked those in the field, particularly during the morning and evening; those in conventional growth chamber conditions did not. Further analysis revealed that fluctuation of irradiance affects transcriptome dynamics in the morning and evening, while fluctuation of temperature only affects transcriptome dynamics in the morning. We found upregulation of genes related to biotic and abiotic stress, whose expression was affected by environmental factors that cannot be mimicked by SmartGC. Our results accelerate the understanding of plant responses to field environments for both field and laboratory studies.

---

To optimise agricultural crop productivity and understand plant behaviour in natural environments, knowledge of plant responses to fluctuating field environments is essential. Numerous studies conducted in controlled environments, such as growth chambers and greenhouses, have facilitated the understanding of plant responses to environmental stimuli. However, such responses are sometimes different from those in controlled environments^1–7^ due to differences between the two environments. Field environments experience daily fluctuations and gradual changes, particularly around dawn and dusk, whereas controlled environments usually fluctuate quickly and regularly between fixed (i.e. square-wave) conditions, which are constant during the day and night, and abruptly transition at dawn and dusk. Light quality, such as red light to far-red light ratio and the presence of ultraviolet-B light, also varied between field and controlled environments. Additionally, plants in the field experience abiotic and biotic stresses, such as wind, precipitation, and insect and pathogen attacks. Such factors make it difficult to apply knowledge obtained from laboratory studies to the field studies in plant science.

To reveal plant responses to fluctuating field environments, field and laboratory studies have attempted to address the differences between the two settings. One approach involves transcriptome analysis of field-grown plants^2, 3, 8–12^. We previously developed a statistical model that predicts the transcriptome dynamics of rice leaves in the field using meteorological data^2^. The modelling approach provides valuable information about plant responses to the field environment, although the detailed mechanism still requires examination under laboratory conditions. Another approach is to mimic the field environment using laboratory equipment. Studies using this approach have clarified photosynthesis characteristics under fluctuating light^6, 13–19^, successfully mimicked primary metabolism of Arabidopsis leaves^4, 5^, and determined the expression patterns of the Arabidopsis florigen gene, *FLOWERING LOCUS T* (*FT*)^7^, in field environments.

Previous studies have clarified the characteristics of plant responses to fluctuating field environments. However, a comprehensive understanding of the differences between plants grown in field and controlled environments is still lacking. Therefore, we developed SmartGC, a high-performance growth chamber that can reproduce fluctuating field environments to compare plant responses to field and controlled environments. By analysing massive transcriptome data of rice plants grown under field and SmartGC conditions, we revealed fillable and unfillable gaps in plant responses to field and controlled environments.

## Results

### Reproduction of environmental field conditions with SmartGC

SmartGC can control irradiance with a 1-second resolution and temperature and relative humidity with a 1-minute resolution (Supplementary Fig. 1), enabling the reproduction of fluctuating field environments. We grew rice plants in SmartGC for 17 days under square-wave conditions. Subsequently, we transferred the plants to five different treatment conditions, where they were left to acclimate for two days. The leaves were sampled on the third day (Fig. 1a). First, we conducted experiments in the field (FIELD) to measure irradiance (light, L), temperature (T), and relative humidity (H) (Supplementary Fig. 1). We then conducted an experiment in SmartGC simulating the environmental factors of FIELD (fluctuating L, T, and H; FL/FTH). A square-wave condition experiment was also conducted using SmartGC (constant L, T, and H; CL/CTH). To distinguish the effect of environmental factors on rice transcriptomes, we also set conditions where only irradiance (FL/CTH) or temperature and humidity (CL/FTH) were fluctuating, while the other factors were held constant. SmartGC successfully simulated irradiance, temperature, and humidity fluctuations for FIELD, except for simulations with high irradiance and high or low humidity due to the setting limitations of the growth chamber (Fig. 1b–c, Supplementary Fig. 2–4).

**Fig. 1.**
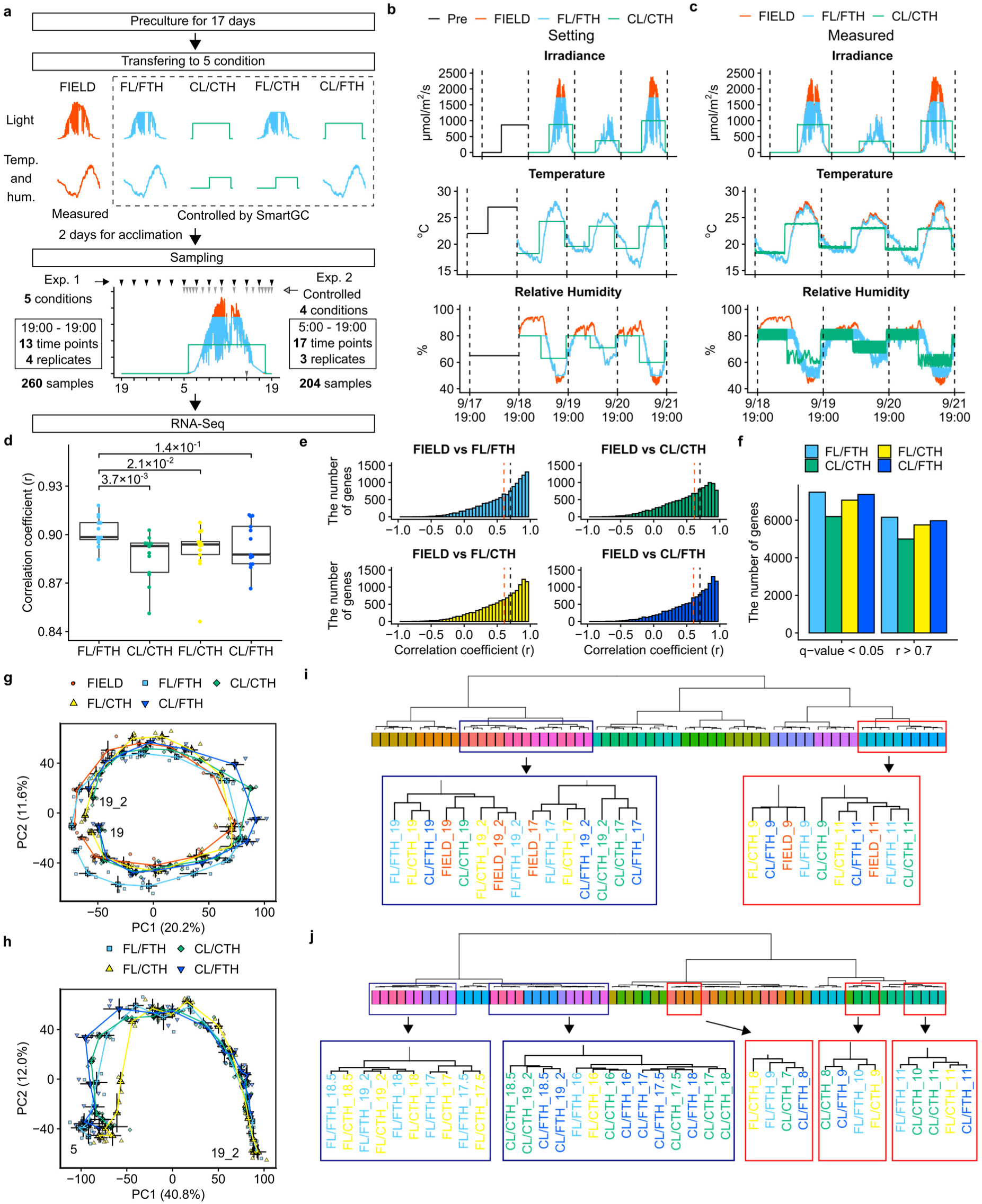
Transcriptome dynamics of rice leaves in the field are mimicked under simulated field environments in SmartGC. **a**, Experimental design of this study. **b**, Irradiance, air temperature, and relative humidity in the preculture, FIELD, FL/FTH, and CL/CTH conditions. The unit of irradiance is photon flux density (PFD) defined over 380–780 nm. **c**, Measured values of irradiance, air temperature, and relative humidity in FIELD, FL/FTH, and CL/CTH. **d**, Boxplots showing pairwise Pearson’s correlation coefficients (r) of transcriptomes between FIELD and the other conditions at each time point. Adjusted p-values of Wilcoxson rank sum test between FIELD vs. FL/FTH and FIELD vs. CL/CTH, FL/CTH, and CL/FTH are shown. Each point shows the mean value of four replicates. **e**, Histogram of pairwise Pearson’s correlation coefficient of expression levels of each gene between FIELD and the other conditions. **f,** A bar graph showing the number of genes with r > 0.7 and q-value < 0.05 in (**e**). **g**, **h**, Principal component analysis (PCA) of transcriptomes in (**g**) Experiment_1 and (**h**) Experiment_2. The percentages of total variance represented by principal component 1 (PC1) and principal component 2 (PC2) are shown in parentheses. Each point shows the mean value of the four replicates; error bars indicate the standard errors of PC1 and PC2. Points were connected by lines according to the time period at each condition. Numbers within the frame indicate the start and end times. 19_2 indicates the timepoint 24 h after the start of sampling day at 19:00. **i**, **j**, Hierarchical cluster dendrograms of the transcriptomes in each condition and time point in (**i**) Experiment_1 (n = 4) and (**j**) Experiment_2 (n = 3). The same-coloured panels in the dendrograms indicate the same sampling time.

### Evaluation of transcriptome similarity between field and controlled environments

We sampled rice leaves from four biological replicates in each condition every 2 h for 24 h (260 total samples) and conducted RNA-Seq analysis (Experiment_1; Fig. 1a, Supplementary Fig. 3–5, Supplementary Table 1). Another experiment was conducted under four controlled conditions to investigate responses around dawn and dusk (Experiment_2; 204 samples obtained from 17 time points [5:00 to 19:00] with three biological replicates). The differences in rice plant growth among the five conditions were not obvious; therefore, leaves at the same stage were sampled in all conditions.

We evaluated transcriptome similarity between conditions using correlation and principal component analysis (PCA). The pairwise Pearson’s correlation coefficient (r) for each time point and condition tended to be higher for FIELD versus FL/FTH than for FIELD versus the other conditions (Fig. 1d). The number of genes with r > 0.7 and the number of statistically significant genes by correlation (p-value adjusted for multiple comparison test [q-value] < 0.05) were highest for FL/FTH and lowest for CL/CTH (Fig. 1e–f). These results indicate that FL/FTH reproduced the rice transcriptome dynamics of FIELD better than did CL/CTH. PC1 in the Experiment_1 PCA separated samples harvested in the light from those harvested in the dark, and PC2 separated those in the morning from those in the afternoon (Fig. 1g, Supplementary Fig. 6). Consequently, samples were ordered by time, suggesting that the diurnal transcriptome dynamics were similarly formed by circadian changes under the five conditions. Hierarchical clustering based on Pearson’s correlation coefficient separated the samples by sampling time points (Fig. 1i, Supplementary Fig. 7). However, the 9:00 CL/CTH samples were clustered with 11:00 samples of all five conditions, whereas the 9:00 samples of the other conditions were clustered together (Fig. 1i, Supplementary Fig. 7). Furthermore, 19:00_2 (24 h after the start of sampling at 19:00) CL/CTH and CL/FTH samples were clustered with 17:00 samples of all five conditions, whereas the 19:00_2 samples of the other conditions were clustered together (Fig. 1i, Supplemental Fig. 7). These results suggest that the internal time progression of the samples was faster around 9:00 in CL/CTH and slower around 19:00_2 in CL/CTH and CL/FTH compared with the other conditions, reflecting the differences in irradiance. This was also supported by the internal time inference of transcriptome samples using the molecular timetable method^20, 21^ (Supplementary Fig. 8, Supplementary Table 2). These results are consistent with a previous study on statistical modelling with transcriptome data^3^, which found that the internal time progression in conventional growth chamber conditions was faster after lights-on and slower before lights-off than in the field. Experiment_2 samples were also separated by time, excluding those in the morning, using PCA and hierarchical clustering (Fig. 1h,j, Supplementary Fig. 7,9). Samples at 7:00 under CL/CTH were clustered with the 8:00 FL/CTH and CL/FTH samples and the 9:00 FL/FTH samples (Fig. 1j, Supplementary Fig. 7). This time lag between conditions continued until 11:00, suggesting that the morning internal time progression was affected by temperature, humidity, and irradiance.

We then evaluated transcriptome similarity between conditions using differentially expressed gene (DEG) analysis at each sampling time point. We compared FIELD with the other conditions using DEG analysis in Experiment_1 (Supplementary Table 3). In this paragraph, DEG indicates the differentially expressed genes between FIELD and the other conditions. There tended to be fewer DEGs in FL/FTH than in the other conditions (Fig. 2a). This is consistent with the results that the rice transcriptome dynamics in FIELD were better reproduced under FL/FTH than the other conditions (Fig. 1d–f, i). The number of DEGs peaked at 7:00 in CL/CTH and FL/CTH, and at 19:00_2 in CL/CTH and CL/FTH (Fig. 2a). Since temperature and humidity were equal in CL/CTH and FL/CTH, these results suggest that the difference between FIELD and CL/CTH in the morning was mainly due to temperature and/or humidity. In contrast, irradiance was equal for CL/CTH and CL/FTH, suggesting that the differences between FIELD and CL/CTH in the evening were mainly due to irradiance. Unlike at 19:00_2, no clear differences between FIELD and CL/CTH were observed at 19:00 (Fig. 2a). This may reflect the weather differences before sampling: the second day was cloudy, while the third day was sunny (Fig. 1b–c). We investigated the overlap of DEGs at each time point and characterised genes affected by environmental conditions into four types (Fig. 2c,e–g): genes affected by light (LIGHT); genes affected by temperature and humidity (TH); genes affected by light, temperature, and humidity (LTH); and genes whose expression in the field was not reproduced under controlled conditions (UNREP). The number of UNREP genes peaked at 13:00 (Fig. 2c). This might reflect the high irradiance in FIELD, which was not simulated by SmartGC (Fig. 1b–c). The number of TH genes tended to be higher in the morning, peaking at 7:00, while the number of LIGHT genes tended to be higher in the evening, peaking at 19:00_2 (Fig. 2a). Additionally, the number of LTH genes was highest at 7:00 and second highest at 19:00_2 (Fig. 2c), suggesting the importance of morning light conditions.

**Fig. 2.**
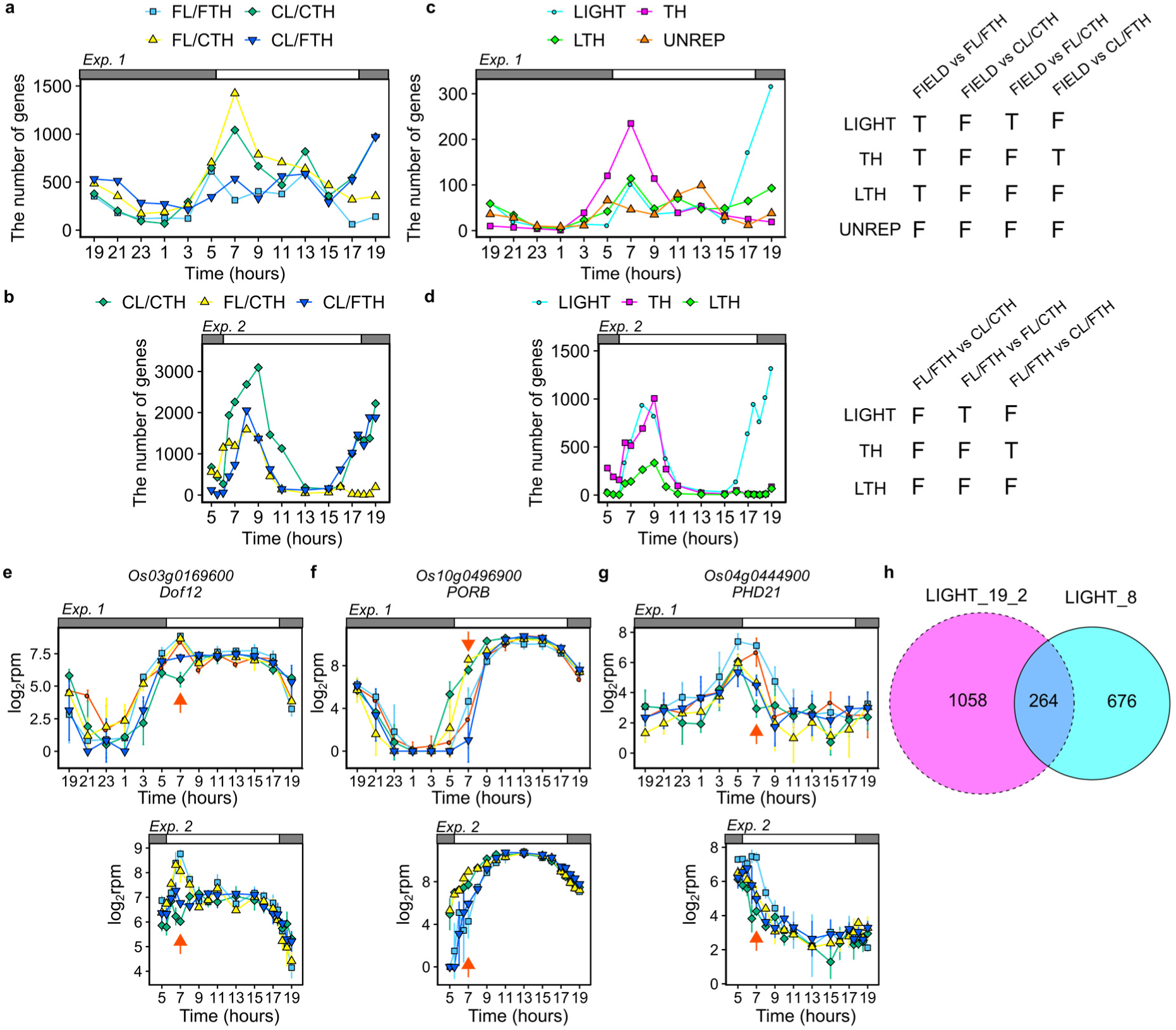
Light affects transcriptome dynamics in the morning and evening while temperature affects them only in the morning. **a**, **b**, The number of DEGs between (**a**) FIELD and the other conditions at each time point in Experiment_1 and (**b**) FL/FTH and the other conditions at each time point in Experiment_2. **c**, **d**, The number of genes affected by environmental conditions at each time point in (**c**) Experiment_1 and (**d**) Experiment_2. Each gene set was selected using the schemes shown in the table. DEGs between conditions that were included and not included in each gene set are shown as T and F, respectively. LIGHT, genes affected by light; TH, genes affected by air temperature; LTH, genes affected by light and air temperature; UNREP, genes whose expressions were regulated by factor(s) other than light and air temperature. **e**–**g**, Expressions of (**e**) LIGHT, (**f**) TH, and (**g**) LTH genes at 7:00 in Experiment_1 and Experiment_2. Points indicate means and error bars indicate standard deviations (n = 4 and n = 3 in Experiment_1 and Experiment_2, respectively). Red arrows indicate 7:00. rpm, reads per million. **h**, Venn diagram of LIGHT genes in the morning (8:00) and evening (19:00_2).

The number of DEGs between FL/FTH and the other conditions in Experiment_2 peaked in the morning and evening for CL/CTH and CL/FTH, and only peaked in the morning for FL/CTH (Fig. 2b, Supplementary Table 4). The overlap of DEGs in Experiment_2 showed that the number of LIGHT genes was high in the morning and evening, while that of TH genes was high only in the evening (Fig. 2d). These results confirm the findings from PCA and hierarchical clustering (Fig. 1g–j). Interestingly, >50% of the LIGHT genes at 8:00 and 19:00_2 overlapped (Fig. 2h), suggesting that the effect of irradiance on transcriptome dynamics was different between the morning and evening. The number of TH genes was higher than that of LTH and LIGHT genes from 5:00 to 6:00 (Fig. 2d), indicating that temperature and humidity began affecting the transcriptome before dawn. In contrast, the number of LTH and LIGHT genes increased from 6:00 to 6:30, indicating that light began affecting the transcriptome 0.5–1 h after dawn. Since the start of dawn only differed by 10 min between FL/FTH and CL/CTH (Supplementary Fig. 4), the gradual versus sudden increase of irradiance, and not the difference in the timing of dawn, caused the increase of LTH and LIGHT genes after dawn. Likewise, the number of TH genes increased beginning 1 h before dusk (16:00–17:00) (Fig. 2d), indicating that a gradual versus sudden decrease of irradiance, and not the difference in the timing of dusk, caused the increase of TH genes before dusk. Overall, these results suggest that gradual versus sudden changes in irradiance affect transcriptome dynamics in the morning and evening, whereas changes in temperature and/or humidity only affect in the morning. The number of DEGs between FL/FTH and the other conditions in Experiment_2 tended to be higher than that between FIELD and the other conditions in Experiment_1 (Fig. 2a–d). This may have resulted from fewer environmental factors affecting transcriptome in SmartGC than in the field.

Although we could not distinguish the effects of temperature and humidity on the transcriptome, the effect of temperature could be greater than that of humidity^2^. Therefore, we considered the effect of temperature or humidity as the effect of temperature in subsequent discussions.

### Circadian clock genes respond to fluctuating irradiance and temperature

Among the circadian clock genes (Fig. 3, Supplementary Fig. 10–12), *PSEUDO-RESPONSE REGULATOR 1* (*PRR1*) and *PHYTOCLOCK 1* (*PCL1*) expression clearly differed between conditions (Fig. 3a). *PRR1* expression increased from 7:00 to 9:00 in FIELD, FL/FTH, and FL/CTH, and from 5:00 to 7:00 in CL/CTH and CL/FTH. *PCL1* expression increased from 13:00 to 15:00 in FIELD, FL/FTH, and FL/CTH, and from 11:00 to 13:00 in CL/CTH and CL/FTH. These results suggest that the increases in *PRR1* expression in the morning and *PCL1* in the daytime were affected by irradiance. In Arabidopsis, *REVEILLE* (*RVE*) genes are positive regulators of *PRR5*, *TOC1*, and evening complex genes^22–26^. The expression of two *RVE* genes (*Os06g0728700* and *Os02g0680700*) decreased from 15:00 to 17:00 in FIELD, FL/FTH, and FL/CTH, and from 17:00 to 19:00_2 in CL/CTH and CL/FTH, suggesting the involvement of irradiance in their regulation (Fig. 3b, Supplementary Fig. 12). These results, paired with the finding that irradiance affects *PRR1* and *PCL1* expression (Fig. 3a), suggest that positive regulation of *TOC1* and *LUX* by *RVE* is conserved in rice^27^. In contrast, expression of the other two *RVE* genes (*Os04g0538900* and *Os02g0685200*) decreased from 7:00 to 9:00 in FIELD, FL/FTH, and CL/FTH, and from 5:00 to 7:00 in CL/CTH and FL/FTH (Fig. 3b, Supplementary Fig. 12), suggesting that their expression was affected by diurnal temperature changes. The circadian rhythm of chloroplast gene expression is regulated by nuclear-encoded genes, such as sigma factor *SIG5*^28^. *SIG*5 expression was lower in FIELD, FL/FTH, and FL/CTH than in CL/CTH and CL/FTH at 17:00 and 19:00_2, suggesting that chloroplast gene expression responds to the diurnal irradiance changes via *SIG5* (Fig. 3c). Overall, these results identified circadian clock genes that respond to fluctuating irradiance and temperature. Future studies should clarify the role of individual genes in response to fluctuating environmental conditions.

**Fig. 3.**
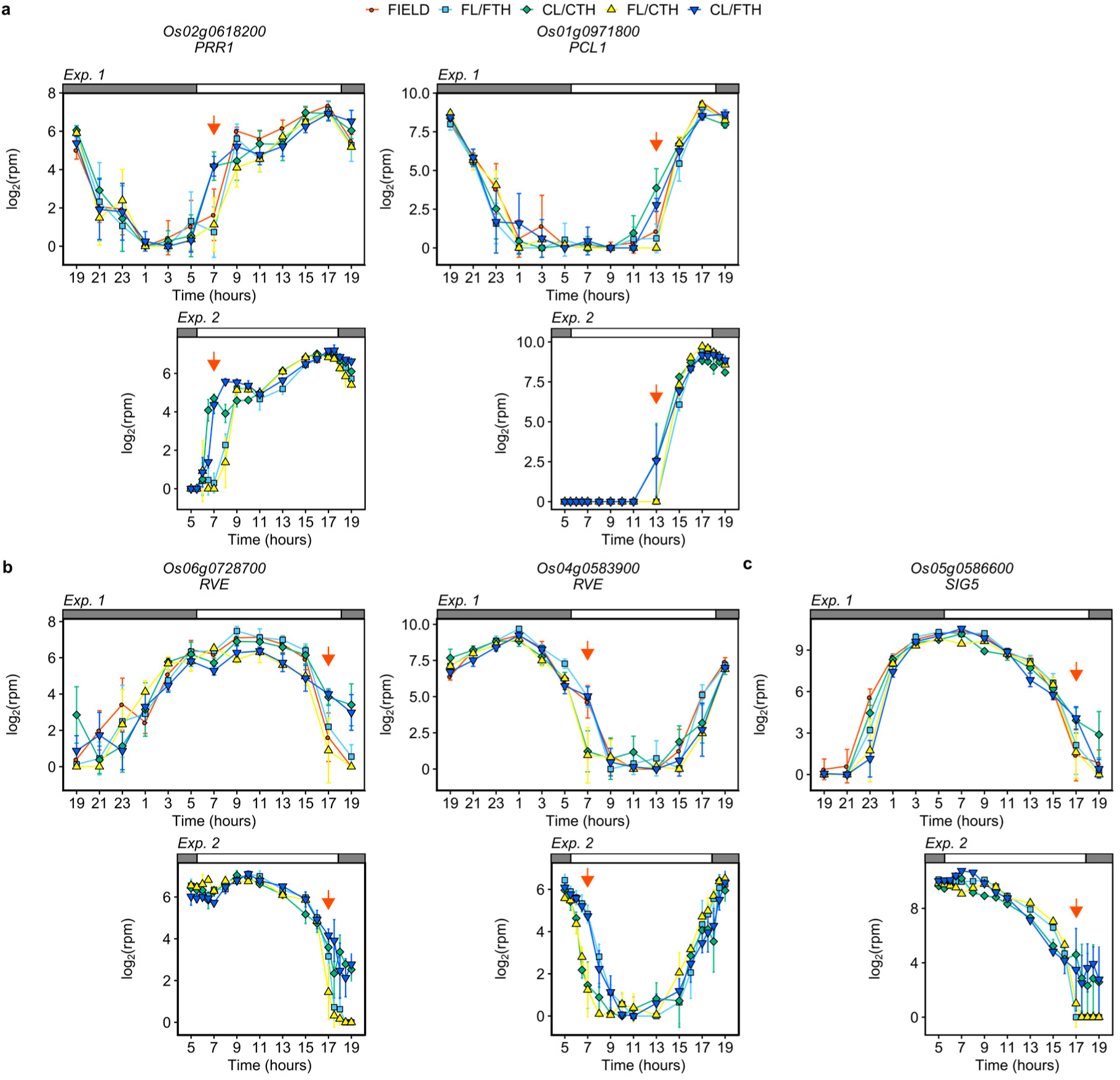
Circadian clocks under field and controlled conditions. Expressions of (**a**) *PSEUDO-RESPONSE REGULATOR 1* (*PRR1*), *PHYTOCLOCK 1* (*PCL1*), (**b**) genes encoding *REVEILLE*, and (**c**) genes encoding sigma factor *SIG5* in Experiment_1 and Experiment_2. Points indicate means and error bars indicate standard deviations (n = 4 and n = 3 in Experiment_1 and Experiment_2, respectively). Red arrows indicate the sampling times discussed in the manuscript. rpm, reads per million.

### Diurnal fluctuation of irradiance affects rice leaf sugar metabolism

To characterise the environmentally-affected genes, we tested for enrichment of genes with annotations in the DEGs detected above (Fig. 4, Supplementary Fig. 13, Supplementary Table 5–12). A total of 7564 and 2942 genes, which have at least one gene ontology (GO) annotation or which belong to one Kyoto Encyclopaedia of Genes and Genomes (KEGG) pathway, respectively, were used for the enrichment test. Genes annotated for photosynthesis (GO:0015979; KEGG pathway: dosa00195) were significantly enriched in DEGs and LTH genes in Experiment_1 (Fig. 4), suggesting that fluctuations in irradiance and temperature affected photosynthesis-related gene expression. Genes annotated for photosynthesis and photosynthesis-antenna proteins (KEGG pathway: dosa00196) or photosynthetic light harvesting (GO:0009765) were significantly enriched in TH genes at 7:00–9:00 in Experiment_1 and 5:00–6:30 in Experiment_2 (Fig. 4b,d, Supplementary Fig. 13). The expression of some photosynthetic light harvesting genes increased before dusk, and this increase occurred earlier for CL/CTH and FL/CTH than for FIELD, FL/FTH, and CL/FTH (Supplementary Fig. 14). These results indicate that photosynthetic light harvesting-related genes are examples of genes whose expression in the morning is affected by gradual changes and fluctuations in temperature. This is consistent with the gene expression model suggested by Nagano et al.^2^. Among the 15 genes annotated for photosynthesis-antenna proteins or photosynthetic light harvesting, the expression of nine genes was affected by night temperature (Supplementary Fig. 14, Supplementary Table 13).

**Fig. 4.**
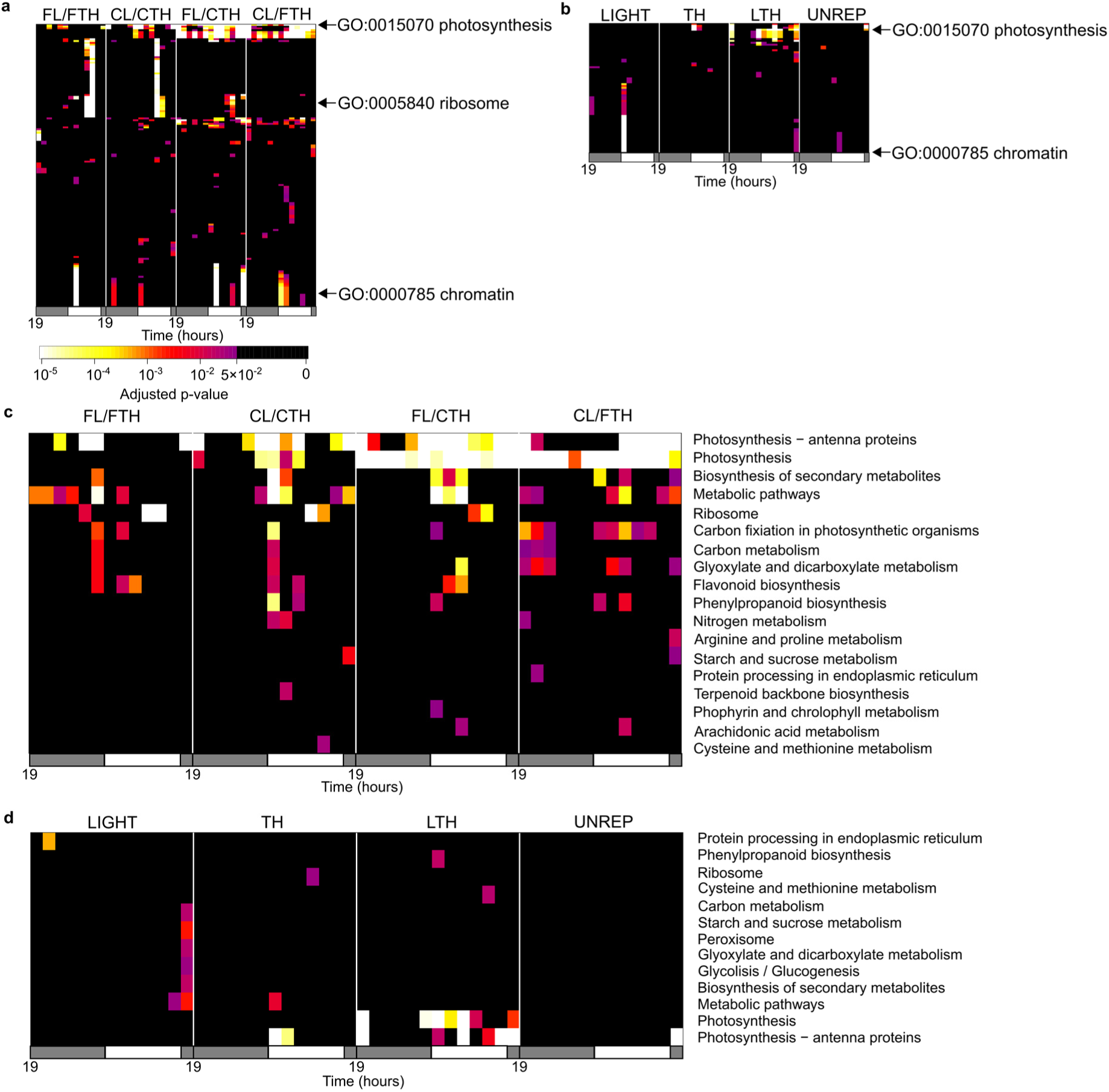
Identification of gene sets affected by fluctuating environmental conditions. Heatmaps of p-values (Fisher’s exact test, two-sided) for significant genes with (**a**, **b**) a particular gene ontology (GO) and (**c**, **d**) a particular KEGG pathway (row) at each time and condition (column) in (**a**, **c**) DEGs between FIELD and the other conditions and (**b**, **d**) LIGHT, TH, LTH, and UNREP genes in Experiment_1. GO and KEGG pathways that have at least one significant (adjusted p-value < 0.05) time point and condition are shown in the heatmaps.

Significant enrichment of sugar metabolism genes was observed in the evening. Genes annotated for starch and sucrose metabolism (KEGG pathway: dosa00500) were significantly enriched in DEGs in FIELD versus CL/CTH and LIGHT genes at 19:00_2 in Experiment_1, and in DEGs in FL/FTH versus CL/CTH from 17:00 to 19:00, FL/FTH versus CL/FTH at 17:30–19:00, and LIGHT genes at 17:30–19:00 in Experiment_2 (Fig. 4c–d, Supplementary Fig. 13). These are examples of genes whose evening expression is affected by fluctuations in irradiance. In Arabidopsis, differences in irradiance between sinusoidal and square-wave conditions affect diurnal changes in carbohydrate content^4, 5^. To clarify the effect of environmental conditions on rice leaf carbohydrate metabolism, we measured the starch and sugar contents in Experiment_1 leaves. Carbohydrate content, especially of sucrose, reflected the differences in diurnal changes of irradiance between conditions (Fig. 1b–c, 5a–b, Supplementary Table 14). In CL/CTH and CL/FTH, starch and sucrose content decreased from dusk to dawn and then increased from dawn to dusk. For sucrose content, a delayed increase at dawn and an early decrease before dusk were observed in FIELD, FL/FTH, and FL/CTH, which is consistent with previous results on Arabidopsis^4, 5^. The delayed increase at dawn was also observed for Arabidopsis leaf starch under sinusoidal conditions with irradiance, but this trend was less prominent in rice. This reflects leaf carbohydrate composition; rice mainly stores sucrose, whereas Arabidopsis mainly stores starch^29^. Although the diurnal trends of changes in starch and sucrose content were similar between FIELD, FL/FTH, and CL/FTH, starch and sucrose content tended to be higher in FIELD than in FL/FTH and FL/CTH. This may have been due to the inability of SmartGC to simulate high irradiance during sunny daytime (Fig. 1b–c) or the difference in the light source (sunlight in FIELD versus LED in SmartGC).

**Fig. 5.**
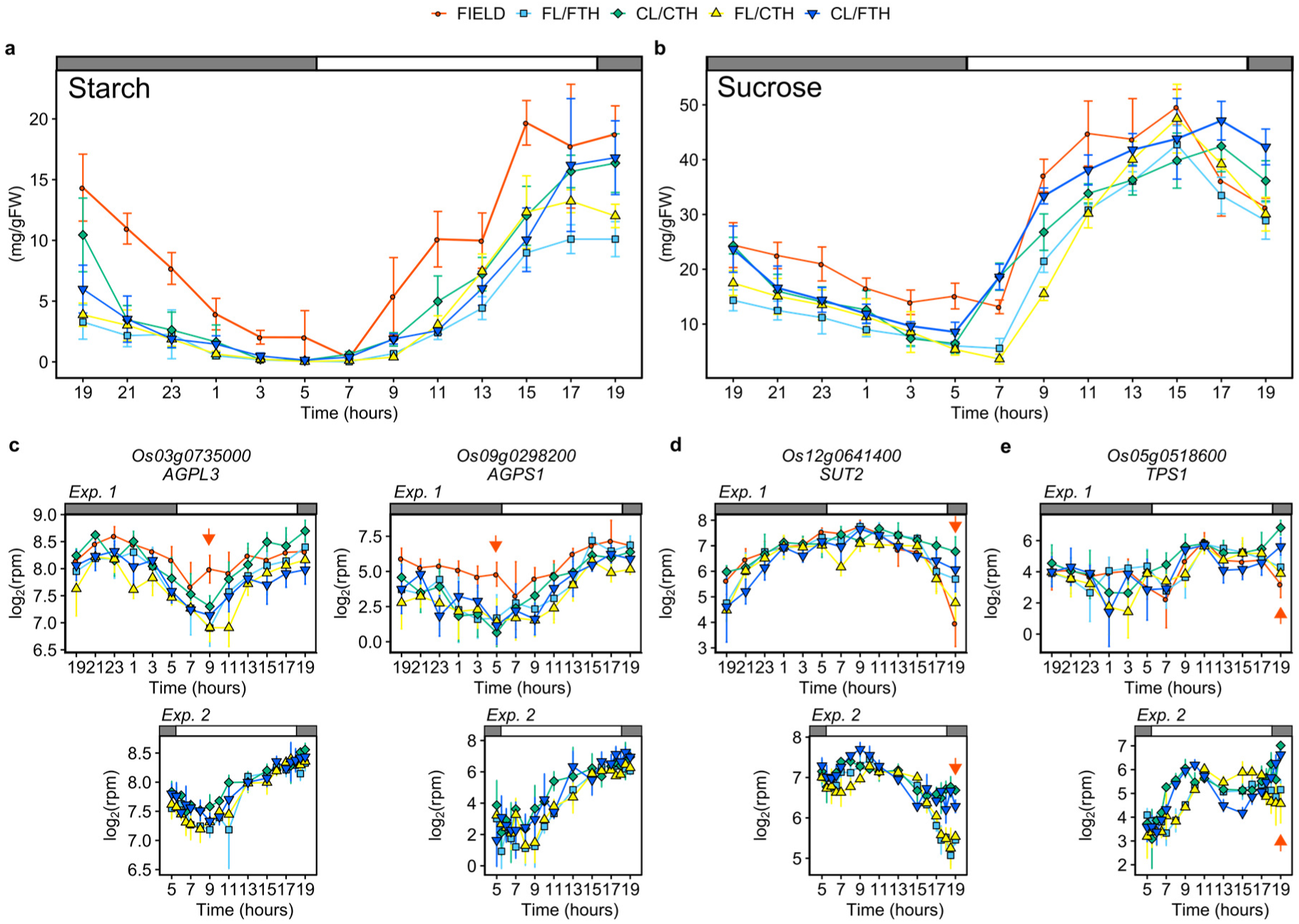
Coordination of leaf sugar metabolism and diurnal change in irradiance. **a**, **b**, Diurnal change in (**a**) starch and (**b**) sucrose content in leaves of each condition in Experiment_1. Results of multiple comparison analysis between conditions using the Tukey– Kramer method are shown in Supplementary Table 14. **c**–**e,** Expressions of genes encoding (**c**) large and small subunits of adenosine diphosphate-glucose pyrophosphorylase (*AGPL3* and *AGPS1*, respectively), (**d**) tonoplast-localised sucrose transporter (*SUT2*), and (**e**) trehalose phosphate synthase (*TPS1*) in Experiment_1 and Experiment_2. Points indicate means and error bars indicate standard deviations (n = 4 and n = 3 in Experiment_1 and Experiment_2, respectively). Red arrows indicate the sampling times discussed in the manuscript. rpm, reads per million. rpm, reads per million. FW, Fresh weight.

Accordingly, with the differences in leaf carbohydrate content, sugar metabolism genes clearly differed between the conditions. Expression of *AGPL3* and *AGPS1*, which encode adenosine diphosphate-glucose pyrophosphorylase (AGP), a key enzyme in starch synthesis^29, 30^, tended to be higher in FIELD than in the other conditions (Fig. 5c). Evening expression of other starch synthesis genes, such as starch synthases (*SSI*, *SSIIb*, and *SSIIIb*) and granule-bound starch synthase (*GBSSII*), showed similar trends to the changes in irradiance; expression decreased earlier in FIELD, FL/FTH, and FL/CTH than in CL/CTH and CL/FTH (Supplemental Fig. 15). This trend was also observed for sugar metabolism and signalling-related gene expression (Fig. 5d–e, Supplementary Fig. 15). For example, expression of sugar transporter (*SUT2*), which regulates carbon export from source leaves to sink organs in rice^31^, showed a similar trend (Fig. 5d). Furthermore, genes encoding trehalose 6-phosphate synthase and trehalose 6-phosphate phosphatase, which belong to the trehalose biosynthesis pathway and play a significant role in sugar signalling^32, 33^, showed similar trends to the leaf sucrose content (Fig. 5e, Supplementary Fig. 16). Overall, these results indicate that differences in diurnal changes in irradiance between FIELD and CL/CTH conditions affected the carbon status and expression of sugar metabolism genes in rice leaves, especially in the evening.

### Field-specific expression of genes related to biotic and abiotic stress

Gene enrichment tests for DEGs showed gene expression specific to FIELD. Genes annotated for ribosomes (GO:0005840; KEGG pathway: dosa03010) were significantly enriched in DEGs in FIELD versus FL/FTH, CL/CTH, and FL/CTH at 13:00 and 15:00 (Fig. 4a). Ribosome-annotated gene expression was significantly higher in FIELD than in the other conditions (Fig. 6a). These results suggest that ribosome-related gene expression was upregulated by unevaluated environmental factor(s) that differed between field and SmartGC experiments. Considering that ribosomal gene expression responds to various biotic and abiotic stresses^34^, the upregulation of ribosome-related genes was probably a response to stressors specific to the field environment. Genes annotated for chromatin (GO:0000785) were significantly enriched in DEGs between FIELD and the other conditions, especially in the morning, indicating the difference in chromatin state dynamics between sunlight and LEDs (Fig. 4a–b, Supplementary Fig. 17).

**Fig. 6.**
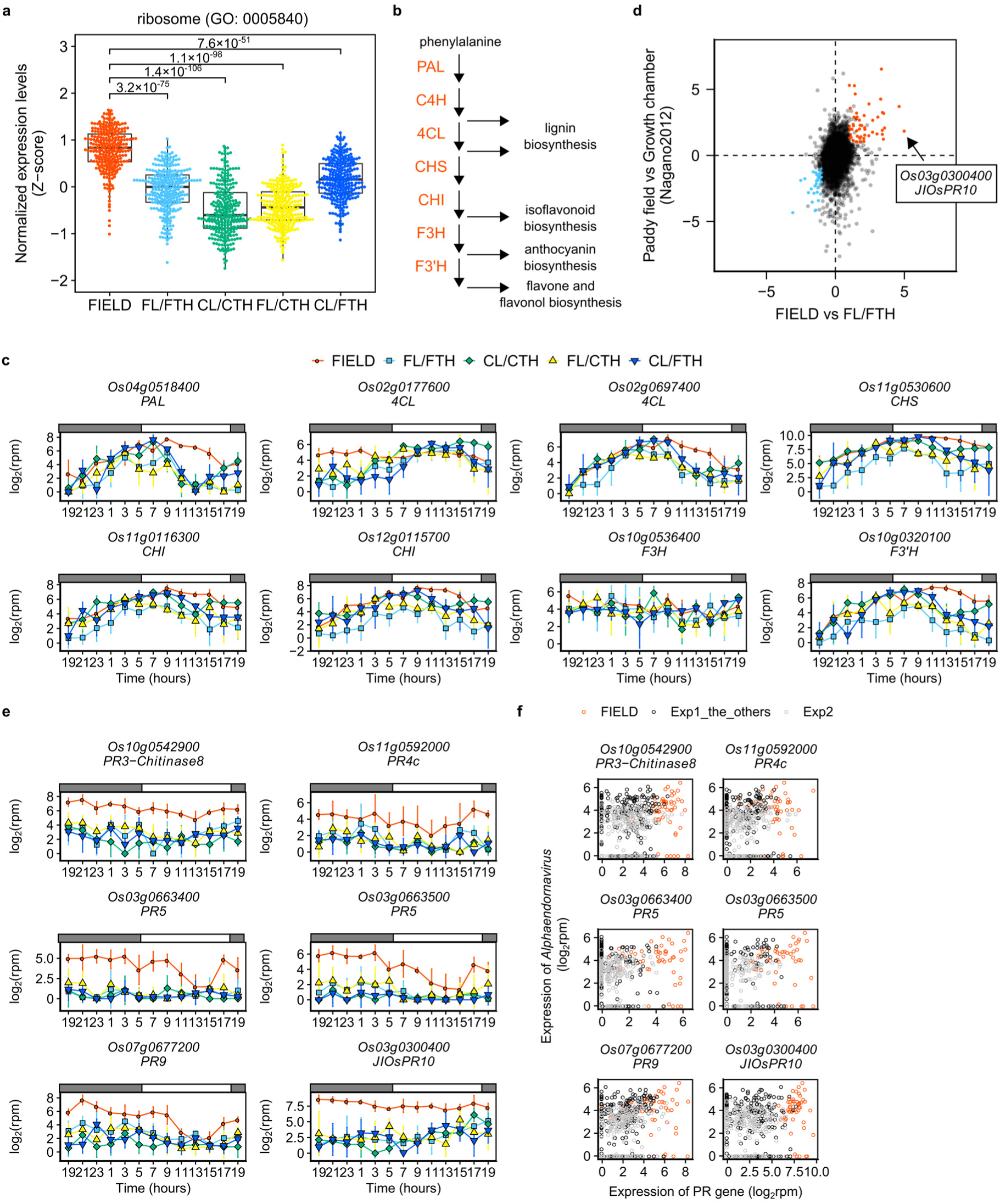
Field-specific expression of genes related to biotic and abiotic stress. **a**, Expression of ribosome-related genes is upregulated in FIELD. Boxplot showing the normalised expression levels (z-score) of genes with annotations for ribosomes (GO:0005840) between FIELD and the other conditions at 13:00 in Experiment 1. Adjusted p-values of Wilcoxon rank sum test between FIELD and the other conditions are shown. **b**, Outline of the phenylpropanoid biosynthesis pathway. Enzymes catalysing each reaction are shown in red. CHI, chalcone isomerase; CHS, chalcone synthase; C4H, cinnamate 4-hydroxylase; F3H, flavanone 3-hydroxylase; F3’H, flavonoid 3-hydroxylase; 4CL, 4-coumarate:coenzyme A ligase; PAL, L-phenylalanine ammonia-lyase. **c**, Expression of genes related to the phenylpropanoid biosynthesis pathway. Points indicate means and error bars indicate standard deviations (n = 4). **d**, Scatter plot showing the differences in the mean expression value between FIELD and FL/FTH in this study and between paddy field and the growth chamber in Nagano et al.^2^. Genes whose mean value of expression was more than 2.0 times higher or lower in both experiments and significantly different between FIELD and FL/FTH at one or more time points are shown as red and blue points, respectively. **e**, Expressions of PR genes whose expressions were upregulated in FIELD. Points indicate means and error bars indicate standard deviations (n = 4). **f,** Scatter plot showing the relationship between the expressions of *Alphaendornavirus*^41^ and PR genes whose expressions were upregulated in FIELD.

Genes annotated for secondary metabolite (KEGG pathway: dosa01110), phenylpropanoid biosynthesis (KEGG pathway: dosa00940), and flavonoid biosynthesis (KEGG pathway: dosa00941) were significantly enriched in DEGs between FIELD and the other conditions, and LTH and UNREP genes (Fig. 4c–d). Phenylpropanoid biosynthesis-related gene expression (Fig. 6b) was significantly upregulated in FIELD compared to the other conditions, mainly from 9:00 to 15:00, except for a gene encoding 4-coumarate:coenzyme A ligase (*Os02g0177600*), which was upregulated at night (Fig. 6c). Since phenylpropanoid biosynthesis-related gene expression is induced by various biotic and abiotic stresses^35, 36^, these results suggest that the upregulation of these genes was a response to field environment stresses.

To investigate field-specific gene expression, we focused on the transcriptome differences between FIELD and FL/FTH. We calculated the mean value of each gene’s expression at all time points and extracted genes whose expression was 2× higher or lower in FIELD than in FL/FTH. We also extracted genes whose expression significantly differed between FIELD and FL/FTH at one or more time points. A total of 159 and 78 genes were identified as upregulated and downregulated, respectively, in FIELD (Fig. 6d, Supplementary Tables 15– 16). Phenylpropanoid biosynthesis-related genes were observed among the highly expressed genes in FIELD (Supplementary Table 15). We also found several genes encoding pathogenesis-related (PR) proteins (Fig. 6e, Supplementary Table 15), which are induced by pathogen attack and are a key component of systemic acquired resistance (SAR)^37^. *NONEXPRESSOR OF PATHOGENESIS-RELATED GENES 1* (*NPR1*) is important for establishing SAR and indirectly activating PR gene expression^37^. However, *NPR1* and the other NPR genes were not upregulated in FIELD (Supplementary Fig. 18), suggesting that PR gene upregulation was independent of NPR genes. Terpene synthesis-related genes, which defend against herbivore- and pathogen-caused tissue damage^38^, were also upregulated in FIELD (Supplementary Table 15, Supplementary Fig. 19). However, we did not observe any signs of herbivory or herbivorous insects. In addition, herbivory-induced early defence signalling genes^39^ were neither upregulated nor downregulated, except for *WRKY30* downregulation in FIELD (Supplementary Fig. 20). Therefore, PR and terpene synthesis-related gene upregulation were likely independent of the effect of insects.

Although no pathogen infection symptoms were observed, upregulation of PR and terpene synthesis-related genes may have resulted from pathogen infection. Therefore, we attempted to detect viral and fungal infections from RNA-Seq data using our previously reported pipeline^40^ (Supplementary Fig. 21) and de novo assembly of unmapped reads to the rice reference genome (Supplementary Fig. 22). The number of reads of viruses^41^ and fungi with poly(A) tails was not specific for FIELD nor correlated with PR gene expression (Fig. 6f, Supplementary Fig. 21–23, Supplementary Table 17). Although we cannot exclude the possibility of infection by bacteria or viruses without poly(A) tails, these results suggest that upregulation of PR and terpene synthase genes in FIELD was a response to physical environmental factor(s) specific to the field.

To determine whether field-specific gene expression is also present in paddy field rice, we reanalysed the microarray data of rice leaves sampled from a paddy field and a growth chamber, which were previously analysed by Nagano et al.^2^. Up- or downregulation of genes in FIELD were consistent with their study^2^ (Fig. 6d, Supplementary Tables 18–19), suggesting that the field-specific gene expression information we obtained is also applicable to paddy field rice.

## Discussion

Although differences between plants grown in the field and controlled environments are well known^1^, few studies have explored the underlying molecular mechanisms. Here, we established an experimental scheme for using laboratory equipment to evaluate plant responses to fluctuating environments. We revealed diurnal transcriptome dynamics in both environments and their fillable and unfillable gaps. Our results complement those that model plant transcriptome responses to field environments^2, 3^. Gradual changes in irradiance affected transcriptome dynamics in the morning and evening, whereas temperature changes only had an effect in the morning (Fig. 2). Accordingly, our statistical model suggested that the number of genes whose expression was affected by a time-specific temperature was the lowest from noon to dusk^2^. The number of genes whose expression was affected by time-specific daily irradiance was higher in the daytime and highest around noon. There was no difference in the number of genes in the morning and evening. Since the plant circadian clock is dawn-dominant^42–44^, the result that both morning irradiance and temperature affect transcriptome dynamics is likely due to circadian entrainment by irradiance and temperature. Conversely, this suggests that the effect of evening irradiance is independent of circadian regulation. Accordingly, less than a half of the morning and evening LIGHT genes overlapped (Fig. 3f). Since sugar metabolism gene expression corresponded to decreased sucrose content in the evening (Fig. 5, Supplementary Fig. 15–16) and diurnal changes in sugar status affect transcriptome dynamics independent of the circadian clock^43, 45^, the effect of evening irradiance potentially depends on differences in carbon status between conditions.

Field plants experience various biotic and abiotic stresses, such as insect and pathogen attack, wind, and UV light, which were not simulated by SmartGC. We found upregulated genes related to ribosomes, phenylpropanoid biosynthesis, and pathogen defence (Fig. 6), all of which are responses to biotic and abiotic stress^34–36, 46^. This indicates that plants cope with the field environment by upregulating stress-responsive genes, despite the less stressful field environment compared to the stress treatment experiments. Alongside the finding that these genes are also upregulated in paddy fields (Fig. 6f), our study suggests that these genes can be targets for rice productivity improvement in the field.

We demonstrated the utility of SmartGC in understanding plant responses to fluctuating environments, although SmartGC cannot completely reproduce field environments, and unfillable gaps between plants grown in field and SmartGC exist. Modifying our experimental scheme will allow the evaluation of plant responses to environmental factors such as humidity, light quality, and light fluctuation, which are unexamined yet important features of the field environment. SmartGC is especially useful for difficult-to-conduct field experiments, such as experiments using genetically modified plants, radioisotopes, or rare environmental conditions. Moreover, SmartGC can be used to predict the effects of future climate change on plants. Further studies utilising SmartGC would bridge the gap between field and laboratory studies and facilitate a comprehensive understanding of plant responses to field environments.

## Methods

### Plant materials and growth conditions

In this study, we developed SmartGC, a high-performance growth chamber (Supplementary Fig. 1). SmartGC is composed of two parts: a growth chamber (LPH-240SP, Nippon Medical & Chemical Instruments Co., Ltd., Osaka, Japan) for controlling temperature and relative humidity, and a Heliospectra L4A LED light source (Heliospectra, Göteborg, Sweden) for controlling light. Both parts were customised to be controlled simultaneously by one computer scheduled to function more than 24 h. The light source can independently control seven types of LEDs with a 1-second resolution. The output value of each LED can be set to 0 or 1 in increments from 15 to 1000 sv (set value). Temperature and relative humidity were set to 15–45 °C and 50–80%, respectively, at a 1-minute resolution. SmartGC records the temperature and relative humidity every minute. Although the light source can set seven types of LEDs independently, we set the output of all LEDs to the same value for each setting.

A common japonica rice *(Oryza sativa* L.) cultivar, Nipponbare, was used in all the experiments in this study. Seeds were sterilised in a 2.5% (v/v) sodium hypochlorite solution for 30 min and then soaked in water at 30 °C for 3 d. Germinated seeds were sown in a cell tray filled with nursery soil (N:P_2_O_5_:K_2_O = 0.6:1.2:1.0 g/kg). Plants were grown in SmartGC for 17 days with a 14 h photoperiod and irradiance level of 867 µmol photon m^-2^ s^-1^ (photon flux density [PFD] of 380–780 nm) 30 cm from the light source, which corresponds to the height of the middle part of the rice leaves used for sampling, by setting an output value of 500 sv (Fig. 1a–b, Supplementary Fig. 1). Plants were then transferred to each condition as follows (Fig. 1a):

i. Field condition (FIELD). We chose a site at Ryukoku University, Otsu, Japan (34° 57’ 43.4” N, 135° 56’ 22.6” E) for the experiment (Supplementary Fig. 1). Plants were transferred at 19:00 on 18 September 2017. During the field experiment, temperature, relative humidity, and irradiance were measured every minute (Fig. 1b). Temperature and relative humidity were measured using THMchip thermo-hygrometers (THM10-TH, FUJIFILM Wako Pure Chemical Corporation, Osaka, Japan) which were set in an aspirated radiation shield^47^. Irradiance was measured using a quantum meter (LA-105, Nippon Medical & Chemical Instruments Co., Ltd.). Because we could not obtain irradiance from 16:00 to 17:31 on 18 September 2017 (i.e. 21 h to 22 h and 31 min after transferring plants to the field), we regarded the change in irradiance during this time as a linear decrease and used the calculated the value of irradiance for further experiments (Fig. 1b–c). The daily light integral was 30, 13, and 34 mol photons d^-1^ on the first, second, and third days, respectively.
ii. Fluctuating light, temperature, and humidity (FL/FTH). Fluctuations in irradiance, temperature and relative humidity in the FIELD condition were simulated. Irradiance in the FIELD condition was simulated every minute by translating irradiance to the light source output using a calibration curve (Supplementary Fig. 24). We measured the PFD of the output from 0, 100, 200, 300, 400, 500, 600, 700, 800, 900, 950, and 1000 sv for each LED and constructed a calibration curve using linear regression. Irradiance (PFD) < 1 in the FIELD was regarded as darkness, and the output value of the light source was set to zero. Because the lowest output value for turning on the light source was 15 sv, the output value during the day was set to 15 sv, as the irradiance in the FIELD was lower than the irradiance in SmartGC with an output value of 15 sv (Fig. 1b, Supplementary Fig. 2, 4). In addition, the highest output value for the light source was 1000 sv, and the output value during the day was set to 1000 sv when the irradiance in the FIELD was higher than the irradiance at an output value of 1000 sv in SmartGC (Fig. 1b). Irradiance was measured every minute using a quantum meter (LA-105) without plants (Fig. 1c; Supplementary Fig. 2). The temperature and relative humidity in the FIELD were simulated every 1 min. The relative humidity in the FL/FTH condition was less than 50% or higher than 80%, so the humidity was set to 50% or 80% (Fig. 1b). Temperature and relative humidity were logged every minute (Supplementary Fig. 2).
iii. Constant light, temperature, and humidity (CL/CTH). Irradiance, temperature, and relative humidity were kept constant during the day and night. The time of dawn and dusk in Otsu, Shiga was that recorded by the National Astronomical Observatory of Japan. On each day, irradiance, temperature, and relative humidity were set to a constant value during the day and night, which were the average values in the field condition during the day and the night, respectively. Because we could not obtain the field data 72 h after sampling, the temperature and relative humidity after dusk (17:55 to 19:00) on the third day was set to the value at 19:00 in FIELD. The relative humidity in FIELD was less than 50% or higher than 80%, so the humidity was set to 50% or 80% (Fig. 1b). Irradiance was measured every minute using a quantum meter (LA-105) without plants (Fig. 1c; Supplementary Fig. 4). Temperature and relative humidity were logged every minute (Supplementary Fig. 4).
iv. Fluctuating light with constant temperature and humidity (FL/CTH). Light was set to the values in the FL/FTH condition, and the temperature and relative humidity were set to the values in the CL/CTH condition.
v. Constant light with fluctuating temperature and humidity (CL/FTH). Light was set to the same values as in the CL/CTH condition, and the temperature and relative humidity were set to the values of the FL/FTH condition.

After transferring the plants to each condition, they were acclimatised for 48 h. Sampling was conducted starting at 19:00 every 2 h for 24 h (13 times in total, Experiment_1) (Fig. 1a, Supplementary Fig. 3–5, Supplementary Table 1). Under each condition, the uppermost, fully expanded leaves (the fifth leaves) were sampled from all four plants per sampling point, frozen in liquid nitrogen, and stored at −80 °C for future use. At each sampling point, sampling was completed within five minutes.

Another sampling was conducted under FL/FTH, CL/CTH, FL/CTH, and CL/FTH conditions to investigate plant responses around dawn and dusk in detail (Experiment_2). The experimental scheme was the same as that described above, except the sampling time points and the number of samples differed. Fifty-eight to seventy-two hours after transferring the plants to each of the conditions, three plants were sampled at each of the following time points: 5:00, 5:30, 6:00, 6:30, 7:00, 8:00, 9:00, 10:00, 11:00, 13:00, 15:00, 16:00, 17:00, 17:30, 18:00, 18:30, and 19:00 (17 times in total) (Fig. 1a, Supplementary Fig. 3–5, Supplementary Table 1).

### RNA-Seq analysis

The leaf samples were ground under cryogenic conditions using a Multi-Beads Shocker (Yasui Kikai, Osaka, Japan). Total RNA was extracted using the Maxwell 16 LEV Plant RNA Kit (Promega, Madison, WI, USA). RNA concentration was measured using the broad-range Quant-iT RNA Assay Kit (Thermo Fisher Scientific, Waltham, MA, USA). RNA (500 ng) was used as the input of each sample for library preparation. Library preparation for RNA-sequencing was conducted using Lasy-Seq^48^ version 0.9 or 1.0 (https://sites.google.com/view/lasy-seq/; Supplementary Fig. 5). The library was sequenced using HiSeq 2500 (Illumina, San Diego, CA, USA) at Macrogen (Seoul, South Korea) or Takara (Kusatsu, Japan) with single-end sequencing lengths of 50 bp or 100 bp, respectively. All obtained reads were trimmed using Trimmomatic^49^ (version 0.3.3) using the following parameters: TOPHRED33, ILLUMINACLIP:TruSeq3-SE.fa:2:30:10, LEADING:19, TRAILING:19, SLIDINGWINDOW:30:20, AVGQUAL:20, MINLEN:40, which indicated that reads with more than 39 nucleotides and average quality scores over 19 were reported. Then, the trimmed reads were mapped onto the reference sequences of the IRGSP-1.0_transcript^50^ and the virus reference sequences, which were composed of complete genome sequences of 7457 viruses obtained from NCBI GenBank^10^ using RSEM^51^ (version 1.3.0) and Bowtie^52^ (version 1.1.2) with default parameters.

The reads per million (rpm) were calculated using the nuclear-encoded gene raw count data, excluding the genes encoding rRNA, as described by Kashima et al.^10^. In Experiments 1 and 2, 0.85–3.35 million and 1.11–4.27 million reads per sample, respectively, were used for calculating rpm (Supplementary Fig. 5). A total of 12,741 genes in which the average number of reads was >10 in all Experiment_1 samples was used for the statistical analysis (Supplementary Fig. 5).

### Inference of internal time using the molecular timetable method

We applied the molecular timetable method^20^ to the transcriptome data of Experiment_1 to infer the internal time of each sample, as described by Higashi et al.^21^. First, we selected time-indicating genes whose expression indicated periodicity and high amplitude. To evaluate the periodicity, we prepared 1440 cosine curves, which had different peaks (0–24 h) measured at 1-minute increments. We fitted the curves to the time-course transcriptome data of FIELD in Experiment_1 (52 total samples) and calculated the correlation coefficient (r) to identify the best-fitting cosine curve (Supplementary Fig. 8). The peak time of the best-fitting curve was estimated as the peak time for each gene and was defined as the molecular peak time. Thus, the molecular peak time was estimated from a single gene, and all the genes were estimated individually. Then, to analyse the amplitude, we calculated the average and standard deviation of each gene expression level. The amplitude value (a) was calculated as the standard deviation divided by the average gene expression level (Supplementary Fig. 8). A total of 143 time-indicating genes was selected according to the cut-off values of r = 0.935 and a = 0.15 (Supplementary Table 2). The molecular peak time of the time-indicating genes was covered throughout the day, which ensured the accurate estimation of internal time (Supplementary Fig. 8).

We normalised the expression level of each time-indicating gene using the z-score, which is defined as the value of the individual expression level minus the average expression level, divided by the standard deviation. We then plotted expression profiles composed of the molecular peak time and the normalised expression level for each sampling time (Supplementary Fig. 8). Finally, the internal time was estimated using a plotted expression profile. We prepared 1440 cosine curves with 1-minute differences from each other and fitted them to the expression profiles. We identified the best-fitting cosine curve, and the corresponding peak time was used to indicate the estimated internal time.

To validate the accuracy of inferring the internal time using the time-indicating genes, we calculated the measurement noise as the standard deviation of the difference between the real and estimated expression of each time-indicating gene. The measurement noise of each gene ranged from 91% to 100% (mean ± standard deviation: 99% ± 1%), indicating that 143 time-indicating genes were sufficient for accurately estimating the internal time^20^.

### Determination of starch and sucrose content

Starch and sucrose content were determined as described by Okamura et al.^30^.

### Analysis of public microarray data

We used the microarray data previously analysed by Nagano et al.^2^. This data was available on the GEO website (https://www.ncbi.nlm.nih.gov/geo/; accession numbers: GSE36777 and GSE36595) and had already been normalised and log-transformed. We used 96 samples of rice (cultivar: Norin 8) grown in paddy fields that were sampled in August 2009, 39–98 days after transplantation. We also used 16 samples of the same rice cultivar grown in a growth chamber, which were sampled at 2:00 and 14:00 at 30, 32, 34, and 36 days after sowing. Details about the samples and conditions in the experiment are described in the study by Nagano et al.^2^. The parameters of the gene expression model in their study were obtained from FiT-DB (https://fitdb.dna.affrc.go.jp/).

### Detection of fungal and viral infection of rice leaves using de novo assembly of unmapped reads to the rice reference transcriptome

Since Lasy-Seq^48^ detects RNA with poly(A) tails, unmapped reads to the rice reference genome can contain RNA of fungi and viruses with poly(A) tails. To clarify whether the rice plants sampled in this study were infected by fungi or viruses, we conducted de novo assembly of unmapped reads to the rice reference genome (Supplementary Fig. 22). Raw sequence reads were merged based on the experiments and conditions of each sample. Reads were then trimmed using Trimmomatic^49^ (version 0.33) with the parameters described above and were then mapped to the rice reference genome using Bowtie2 with default parameters, except setting N = 1. After extracting the unmapped reads and removing the duplicated reads, de novo assembly was conducted using Trinity with default parameters. After removing redundant reads using CD-HIT^53^, 19 contigs were identified. Each contig was annotated using BLASTn for nucleotides^54^ (Supplementary Fig. 23, Supplementary Table 17).

### Statistical analysis

All statistical analyses were performed using R software^55^ version 3.5.3. Specifically, differentially expressed gene (DEG) analysis was conducted using R package TCC^56, 57^ version 1.20.0. Normalisation was conducted using iDEGES/edgeR^58^ with a false discovery rate (FDR) of 0.1, and DEG detection was conducted using edgeR with FDR = 0.05. Gene enrichment tests for GO and Kyoto Encyclopaedia of Genes and Genomes^59^ (KEGG) pathways were conducted using the R package GO.db^60^ version 3.6.0 and KEGG.db^61^ version 3.2.3, respectively, as described by Nagano et al.^9^. The FDR was controlled using Benjamini and Hochberg’s method^62^ with FDR = 0.05. Log_2_(rpm) was calculated as log_2_(rpm + 1). Multiple comparison tests for starch and sucrose contents were conducted using the R package car (3.0.10).

### Data and code availability

The scripts used in this study are available at https://github.com/naganolab/SmartGC_RNA-Seq. All datasets generated and/or used in this study are available in the Sequence Read Archive (SRA) under accession number PRJNA726716.

## Acknowledgements

We thank Nippon Medical & Chemical Instruments Co., Ltd. (Osaka, Japan) and LPixel Inc. (Tokyo, Japan) for their help in the establishment of SmartGC, Kyoko Mogami for technical assistance with the RNA-Seq experiments, and Dynacom Co., Ltd. (Chiba, Japan) for technical assistance with the analysis of RNA-Seq data. This work was supported by JST CREST Grant Number JPMJCR15O2, Japan, awarded to AJN.

## Author contributions

AJN conceived the study. YH conducted the field and SmartGC experiments. YN supported the SmartGC experiments. YH, AT, M. Kamitani, M. Kashima, and YK conducted the RNA-Seq experiments. YH analysed the data. YH and AJN wrote the manuscript with input from all co-authors.

## Supplementary Figures and Tables Legend

**Supplementary Fig. 1. SmartGC chamber and field used in this study. a**, **b**, Image of (**a**) SmartGC and (**b**) the field used in this study. **c**, **d**, Rice grown in (**d**) SmartGC 14 days after sowing and (**d**) in the field 17 days after sowing.

**Supplementary Fig. 2. Comparison of set and measured data of environmental conditions. a**, **b**, Irradiance, **c**, **d**, air temperature, and **e**, **f**, relative humidity of (**a**, **c, e**) FL/FTH and (**b**, **d**, **f**) CL/CTH conditions. Sampling time points are marked with arrows. The fluctuation of irradiance in FIELD was simulated in FL/FTH over three days, with the exception of the high irradiance in the sunny midday on the first and third days due to the upper limit of the light source output in SmartGC. Temperature was controlled by SmartGC in a range within 0.9 °C of the set value for FL/FTH and the values for the FIELD condition. Relative humidity was controlled in a range within 9% of the set value for FL/FTH and the values for the FIELD condition, except for high and low humidity (higher than 80 % and less than 50%) due to the upper and lower setting limit of SmartGC.

**Supplementary Fig. 3. Environmental conditions on the sampling day.** Sampling time points are shown with arrows.

**Supplementary Fig. 4. Environmental conditions in the morning and evening on the sampling day. a**, **b**, Irradiance, **c**, **d**, air temperature, and **e**, **f**, relative humidity from (**a**, **c, e**) 5:00–7:00 and (**b**, **d**, **f**) 17:00–19:00 on the sampling day measured in the FIELD condition and set for the FL/FTH and CL/CTH conditions. Sampling time points are shown with arrows.

**Supplementary Fig. 5. Workflow of RNA-Seq data preprocessing. a**, Histogram of the total read counts for each sample. Since some samples were sequenced using more than one library, reads for each sample were merged after mapping the reads to the transcript references. **b**, Histogram of the mean read count for each gene in Experiment_1. After filtering, 12,741 genes were used for statistical analyses.

**Supplementary Fig. 6. Principal component analysis (PCA) of transcriptomes in Experiment_1. a**, PCA of transcriptomes at each time point and condition, which corresponds to Fig. 1g. Each point represents the mean value of the four replicates, and error bars indicate the standard errors of PC1 and PC2. **b**, PCA of transcriptomes of each sample at each time point and condition. Four replicates at each time point and condition are shown independently. Numbers indicate sampling times. The percentages of the total variance represented by PC1 and PC2 are shown in parentheses. 19_2 indicates the timepoint 24 h after the start of sampling day at 19:00.

**Supplementary Fig. 7. Hierarchical cluster dendrogram of transcriptomes of each condition and time point.** The dendrograms in Fig. 1i and Fig. 1j correspond to **a** and **b**, respectively. Mean values of (**a**) 4 and (**b**) 3 replicates at each time point and condition were used for the analyses. 19_2 indicates the timepoint 24 h after the start of sampling day at 19:00.

**Supplementary Fig. 8. Inference of internal time using the molecular timetable method. a**, Histogram showing the Pearson correlation coefficient (r) between the expression of each gene and its best-fitting cosine curve. The red dashed line indicates the selection threshold (r = 0.935) for time-indicating genes. **b**, Histogram showing the expression amplitude of each gene. The red dashed line indicates the selection threshold (a = 0.15) for time-indicating genes. **c**, Histogram showing the molecular peak times of 143 time-indicating genes. **d**, An example of internal time inference using the molecular timetable method. Normalised expressions of time-indicating genes at the molecular peak time of each gene were fitted to the cosine curve, and the peak time indicates the internal time. **e,** Inference of internal time at each condition in Experiment_1. Inferred time minus real time is shown for each sample (n = 4). Progression of internal time in the evening was slower in CL/CTH and CL/FTH than in FIELD.

**Supplementary Fig. 9. Principal component analysis (PCA) of transcriptomes in Experiment_2. a**, PCA of transcriptomes at each time point and condition, which correspond to Fig. 1h. Each point represents the mean value of three replicates, and error bars indicate standard errors of PC1 and PC2. **b**, PCA of transcriptomes of each sample at each time point and condition. Four replicates at each time point and condition are shown independently. Numbers indicate sampling times. The percentages of total variance represented by PC1 and PC2 are shown in parentheses. 19_2 indicates the timepoint 24 h after the start of sampling day at 19:00.

**Supplementary Fig. 10. Expression of genes encoding core circadian clock genes.** Points indicate means and error bars indicate standard deviations (n = 4 and n = 3 in Experiment_1 and Experiment_2, respectively). *LHY*, *LATE ELONGATED HYPOCOTYL*; *PRR73*, *PSEUDO-RESPONSE REGULATOR 73*; *PRR37*, *PSEUDO-RESPONSE REGULATOR 37*; *PRR95*, *PSEUDO-RESPONSE REGULATOR 95*; *PRR59*, *PSEUDO-RESPONSE REGULATOR 59*; *GI*, *GIGANTIA*; *ELF3*, *EARLY-FLOWERING 3*; *ELF4*, *EARLY-FLOWERING 4*.

**Supplementary Fig. 11. Expression of genes encoding photoreceptors.** Points indicate means and error bars indicate standard deviations (n = 4 and n = 3 in Experiment_1 and Experiment_2, respectively). *CRY*, cryptochrome; *PHOT*, phototropin; *PHY*, phytochrome; *UVR8*, UV-B resistance 8; *ZTL*, *ZEITLUPE*.

**Supplementary Fig. 12**. **Expression of genes encoding (a) REVEILLE (RVE), (b) NIGHT LIGHT-INDUCIBLE AND CLOCK-REGULATED protein (LNK), and** (**c**) **PHYTOCHROME INTERACTING FACTOR-LIKE protein (PIL).** Points indicate means and error bars indicate standard deviations (n = 4 and n = 3 in Experiment_1 and Experiment_2, respectively).

**Supplementary Fig. 13. Significant enrichment of genes with specific annotations in DEGs between FIELD and the other conditions in Experiment_2.** Heatmaps of P-values (Fisher’s exact test, two-sided) for significant genes with (**a**, **b**) a particular gene ontology (GO) and (**c, d**) a particular KEGG pathway (row) in each time and condition (column) in (**a, c**) DEGs between FIELD and the other conditions and (**b, d**) LIGHT, TH, LTH, and UNREP genes. GO and KEGG pathways that have at least one significant (adjusted p-value < 0.05) time point and condition are shown.

**Supplementary Fig. 14. Expressions of genes related to light harvesting.** Expressions of genes with annotation for photosynthesis-antenna protein (KEGG pathway: dosa00196) or photosynthetic light harvesting (GO:0009765) in Experiment_1 and Experiment_2 are shown. Red points on the left side of the gene name indicate that the gene was affected by night temperature in Nagano et al.^2^ (See Manuscript and Supplemental Table 13). Points indicate means and error bars indicate standard deviations (n = 4 and n =3 in Experiment_1 and Experiment_2, respectively).

**Supplementary Fig. 15. Expression of genes related to (a) starch and (b) sucrose metabolism.** Points indicate means and error bars indicate standard deviations (n = 4 and n = 3 in Experiment_1 and Experiment_2, respectively). *CIN1*, cytosolic invertase 1; *GPT2*, glucose-6-phosphate/phosphate translocator 2; *GPT2-3*, glucose-6-phosphate/phosphate translocator 2-3; *GBSSII*, Granule-bound starch synthase II; *NIN4*, neutral invertase 4; *PHOH/PHO2*, starch phosphorylase 2; *SSI*, starch synthase I; *SSIIb*, starch synthase IIb; *SSIIIb*, starch synthase *IIIb*; *TMT2*, tonoplast monosaccharide transporter 2; *UGP1*, UDP- glucose pyrophosphorylase 1.

**Supplementary Fig. 16. Expressions of genes encoding (a) trehalose phosphate synthase (TPS) and (b) trehalose phosphate phosphatase (TPP).** Expressions of *TPS1* are shown in Fig. 5e. *TPS9*, *TPS11*, *TPP3*, *TPP4*, *TPP5*, and *TPP8* are not shown because of their low expressions. Points indicate means and error bars indicate standard deviations (n = 4 and n = 3 in Experiment_1 and Experiment_2, respectively).

**Supplementary Fig. 17. Different timing of upregulation of chromatin-related genes between conditions.** Expressions of genes with annotations for chromatin (GO:0000785) in **(a)** Experiment_1 and **(b)** Experiment_2. Each line indicates the mean value of the expressions of each gene (n = 4 and n = 3 in Experiment_1 and Experiment_2, respectively). Expressions of genes with annotation for chromatin were induced at 5:00–7:00 in CL/CTH and CL/FTH, and at 7:00–9:00 in FL/FTH and FL/CTH, but the induction in the morning was not obvious in FIELD. The timing of induction was reproduced in Experiment_2. These results suggest that the expression of genes with annotation for chromatin was induced by LED light, but not by sunlight, in the morning, and the timing of induction was affected by diurnal changes in irradiance.

**Supplementary Fig. 18. Expressions of (a) pathogenesis-related (PR) genes and (b) NONEXPRESSOR OF PATHOGENESIS-RELATED (NPR) GENES in Experiment_1.** Points indicate means and error bars indicate standard deviations (n = 4).

**Supplementary Fig. 19. Expressions of genes related to terpene synthase in Experiment_1.** Points indicate means and error bars indicate standard deviations (n = 4).

**Supplementary Fig. 20. Expressions of genes involved in early defence signalling to herbivore attack listed in Ye et al.**^39^ **in Experiment_1.** Points indicate means and error bars indicate standard deviations (n = 4). Early defence signalling genes include *Oryza sativa* leucine-rich repeat receptor-like kinase 1 (*OsLRR-RLK1*), mitogen-activated protein kinase (*OsMPK3* and *OsMPK6*), WRKY transcription factors (*OsWRKY13*, *OsWRKY24*, *OsWRKY30*, *OsWRKY33*, *OsWRKY45*, *OsWRKY53*, and *OsWRKY70*) and jasmonate synthesis genes (*OsHI-LOX*, *OsAOS1*, *OsAOC*, *OsOPR3*, and *OsJAR1*).

**Supplementary Fig. 21. Detection of virus infection from RNA-Seq data.** Scatter plot showing the max read number and max coverage of 115 viruses with maximum reads per sample >1. *Alphaendornavirus*, which shows no clear symptoms in rice plants^41^, had the highest reads (maximum 166 reads per sample) and coverage of the reads to the reference (6.3% with depth ≥ 3). Correlations between the read number of the *Alphaendornavirus* and the expressions of PR genes are shown in Fig. 6f. The number of reads and the coverage of the *Alphaendornavirus* were low in comparison with infected samples discussed in Kamitani et al.^40^ (more than 10% in a sample with the lowest coverage). Therefore, this result suggests that even if *Alphaendornavirus* existed in the rice leaves, the copy number of the virus was limited.

**Supplementary Fig. 22. Scheme for searching for genes that were found by de novo transcriptome assembly from unmapped reads to the rice reference transcriptomes.**

**Supplementary Fig. 23. Scatter plot showing the relationship between PR genes and contigs found by de novo transcriptome assembly from unmapped reads to the rice reference transcriptomes.** We selected 6 contigs from 19 contigs after excluding those with annotations for *Alphaendornaviruses*, synthetic constructs, and expression vectors. contig1, exp1_CL/FTH.TRINITY_DN138_c0_g1_i1; contig2, exp1_FL/CTH.TRINITY_DN11_c0_g1_i1; contig3, exp1_FL/CTH.TRINITY_DN249_c0_g1_i2; contig4, exp2_CL/CTH.TRINITY_DN586_c0_g1_i1; contig5, exp2_CL/FTH.TRINITY_DN544_c0_g3_i1; contig6, exp2_FL/CTH.TRINITY_DN119_c0_g1_i1. Details of each gene are shown in Supplementary Table 17.

**Supplementary Fig. 24. Calibration curve of the output of the light source of SmartGC versus irradiance.** Calibration curve for seven types of LED light and the sum of all LEDs are shown. The unit of irradiance is photon flux density (PFD) defined over 380–780 nm.

Supplementary Table 1. Sample attributes used in this study.

Supplementary Table 2. Time-indicating genes used in this study.

Supplementary Table 3. The q-value of DEG analysis between FIELD and other conditions at each time point in Experiment_1.

Supplementary Table 4. The q-value of DEG analysis between FL/FTH and other conditions at each time point in Experiment_2.

Supplementary Table 5. Enriched gene ontology in DEGs between FIELD and other conditions in Experiment_1.

Supplementary Table 6. Enriched gene ontology in LIGHT, TH, LTH, and UNREP genes in Experiment_1.

Supplementary Table 7. Enriched KEGG pathway in DEGs between FIELD and other conditions in Experiment_1.

Supplementary Table 8. Enriched KEGG pathway in LIGHT, TH, LTH, and UNREP genes in Experiment_1.

Supplementary Table 9. Enriched gene ontology in DEGs between FL/FTH and other conditions in Experiment_2.

Supplementary Table 10. Enriched gene ontology in LIGHT, TH, and LTH genes in Experiment_2.

Supplementary Table 11. Enriched KEGG pathway in DEGs between FL/FTH and other conditions in Experiment_2.

Supplementary Table 12. Enriched KEGG pathway in LIGHT, TH, and LTH genes in Experiment_2.

Supplementary Table 13. Parameters of the gene expression model in Nagano et al.^2^ for genes related to photosynthetic light harvesting.

Supplementary Table 14. Multiple comparison tests of starch and sucrose contents between conditions in Experiment_1.

Supplementary Table 15. List of genes whose mean expression was higher in FIELD than in FL/FTH.

Supplementary Table 16. List of genes whose mean expression was lower in FIELD than in FL/FTH.

Supplementary Table 17. List of contigs identified by de novo transcriptome assembly from unmapped reads to the rice reference transcriptome.

Supplementary Table 18. List of genes whose expressions were higher both in paddy fields than in growth chambers and in FIELD than in FL/FTH.

Supplementary Table 19. List of genes whose expression was lower both in the paddy field than in the growth chamber and in FIELD than in FL/FTH.

