## Supplementary Figures1-24 for "Fillable and Unfillable Gaps in Rice Transcriptome under Field and Controlled Environments"

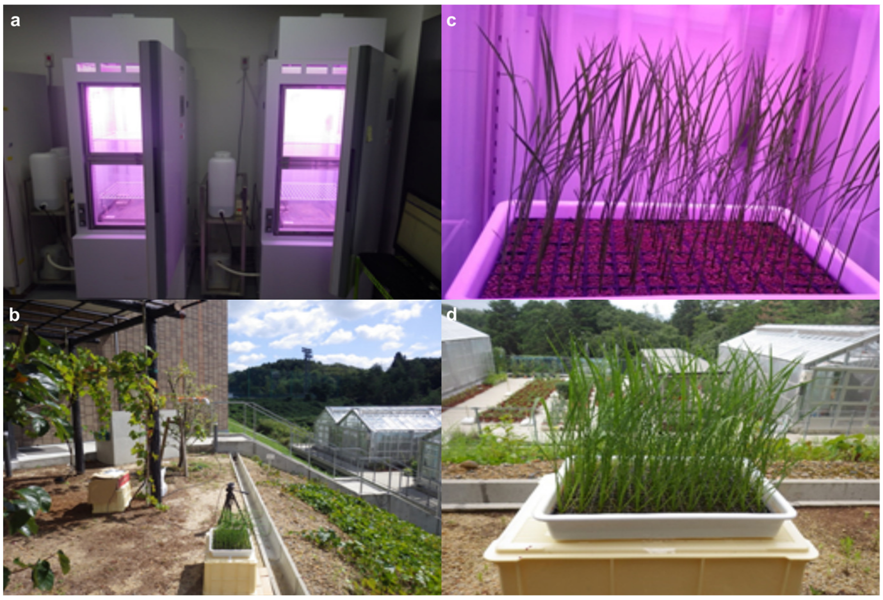


**Supplementary Fig. 1. SmartGC chamber and field used in this study. a**, **b**, Image of (**a**) SmartGC and (**b**) the field used in this study. **c**, **d**, Rice grown in (**d**) SmartGC 14 days after sowing and (**d**) in the field 17 days after sowing.

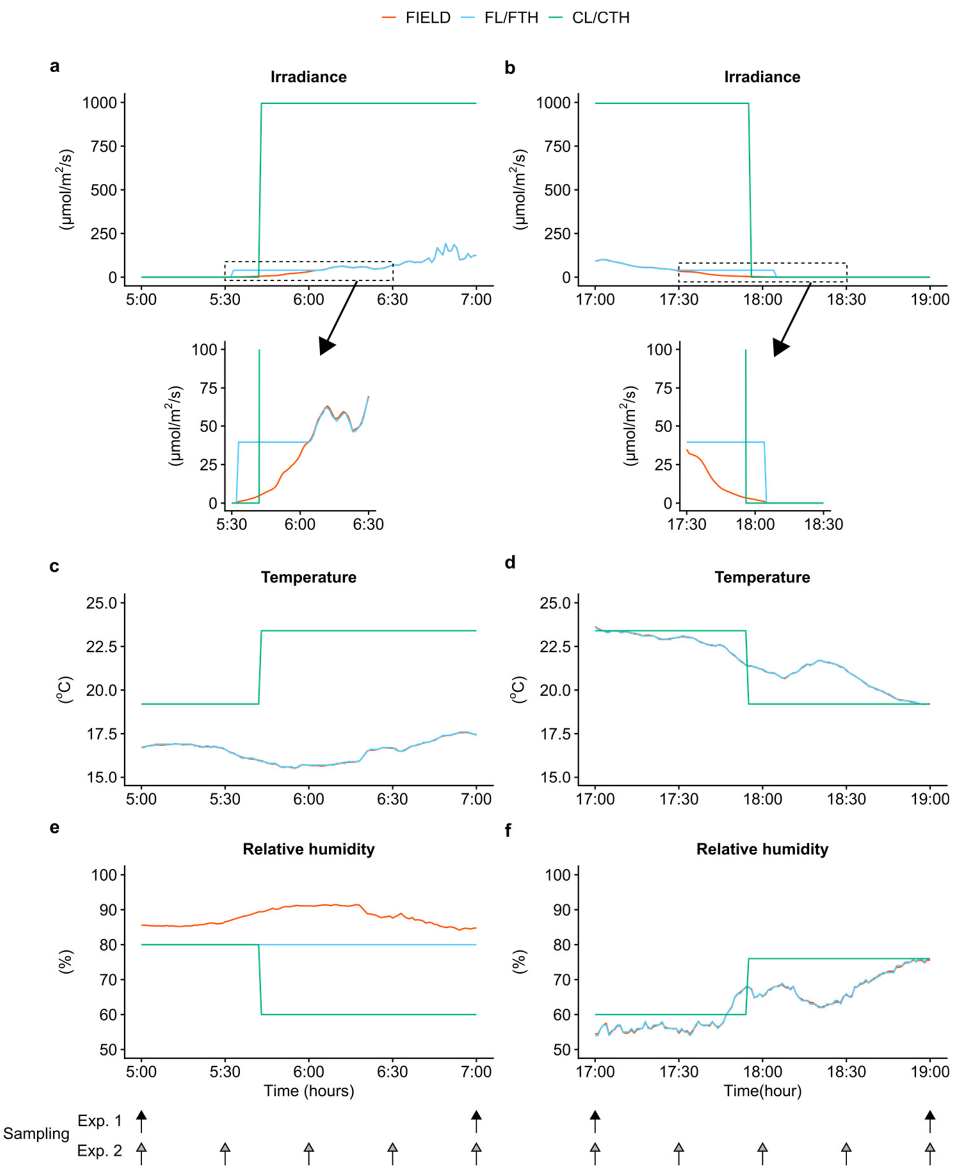


**Supplementary Fig. 2. Comparison of set and measured data of environmental conditions. a**, **b**, Irradiance, **c**, **d**, air temperature, and **e**, **f**, relative humidity of (**a**, **c, e**) FL/FTH and (**b**, **d**, **f**) CL/CTH conditions. Sampling time points are marked with arrows. The fluctuation of irradiance in FIELD was simulated in FL/FTH over three days, with the exception of the high irradiance in the sunny midday on the first and third days due to the upper limit of the light source output in SmartGC. Temperature was controlled by SmartGC in a range within 0.9 °C of the set value for FL/FTH and the values for the FIELD condition.

Relative humidity was controlled in a range within 9% of the set value for FL/FTH and the values for the FIELD condition, except for high and low humidity (higher than 80 % and less than 50%) due to the upper and lower setting limit of SmartGC.

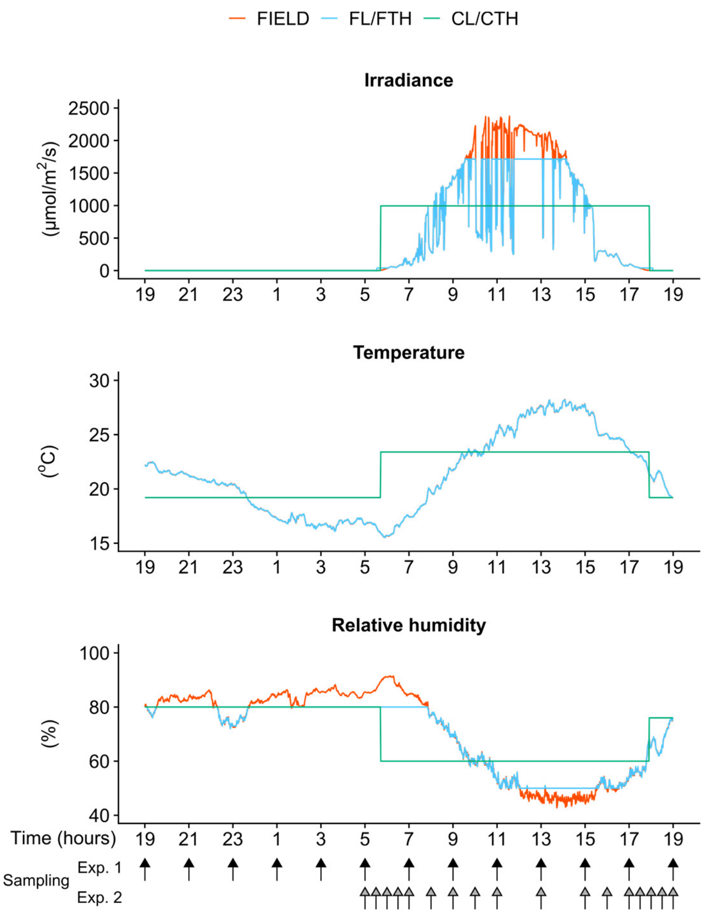


**Supplementary Fig. 3. Environmental conditions on the sampling day.** Sampling time points are shown with arrows.

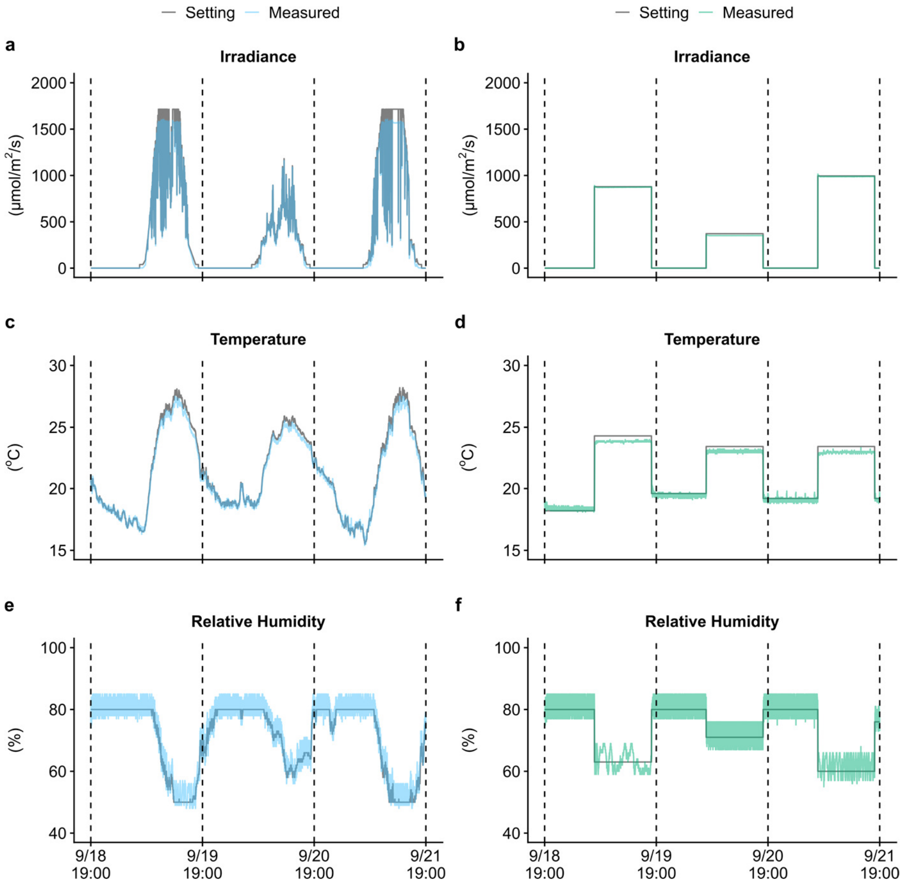


**Supplementary Fig. 4. Environmental conditions in the morning and evening on the sampling day. a**, **b**, Irradiance, **c**, **d**, air temperature, and **e**, **f**, relative humidity from (**a**, **c**, **e**) 5:00–7:00 and (**b**, **d**, **f**) 17:00–19:00 on the sampling day measured in the FIELD condition and set for the FL/FTH and CL/CTH conditions. Sampling time points are shown with arrows.

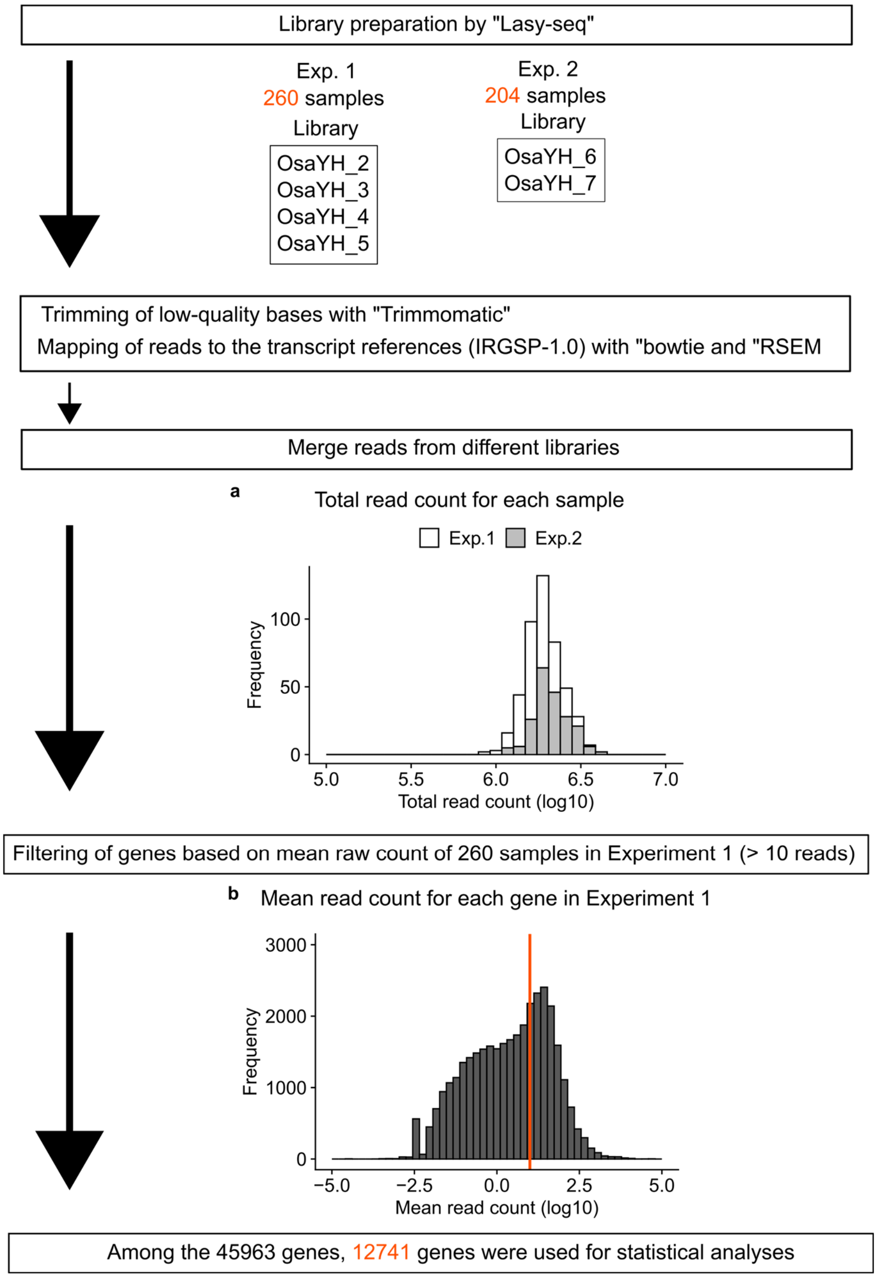


**Supplementary Fig. 5**. **Workflow of RNA-Seq data preprocessing.** **a**, Histogram of the total read counts for each sample. Since some samples were sequenced using more than one library, reads for each sample were merged after mapping the reads to the transcript references. **b**, Histogram of the mean read count for each gene in Experiment_1. After filtering, 12,741 genes were used for statistical analyses.

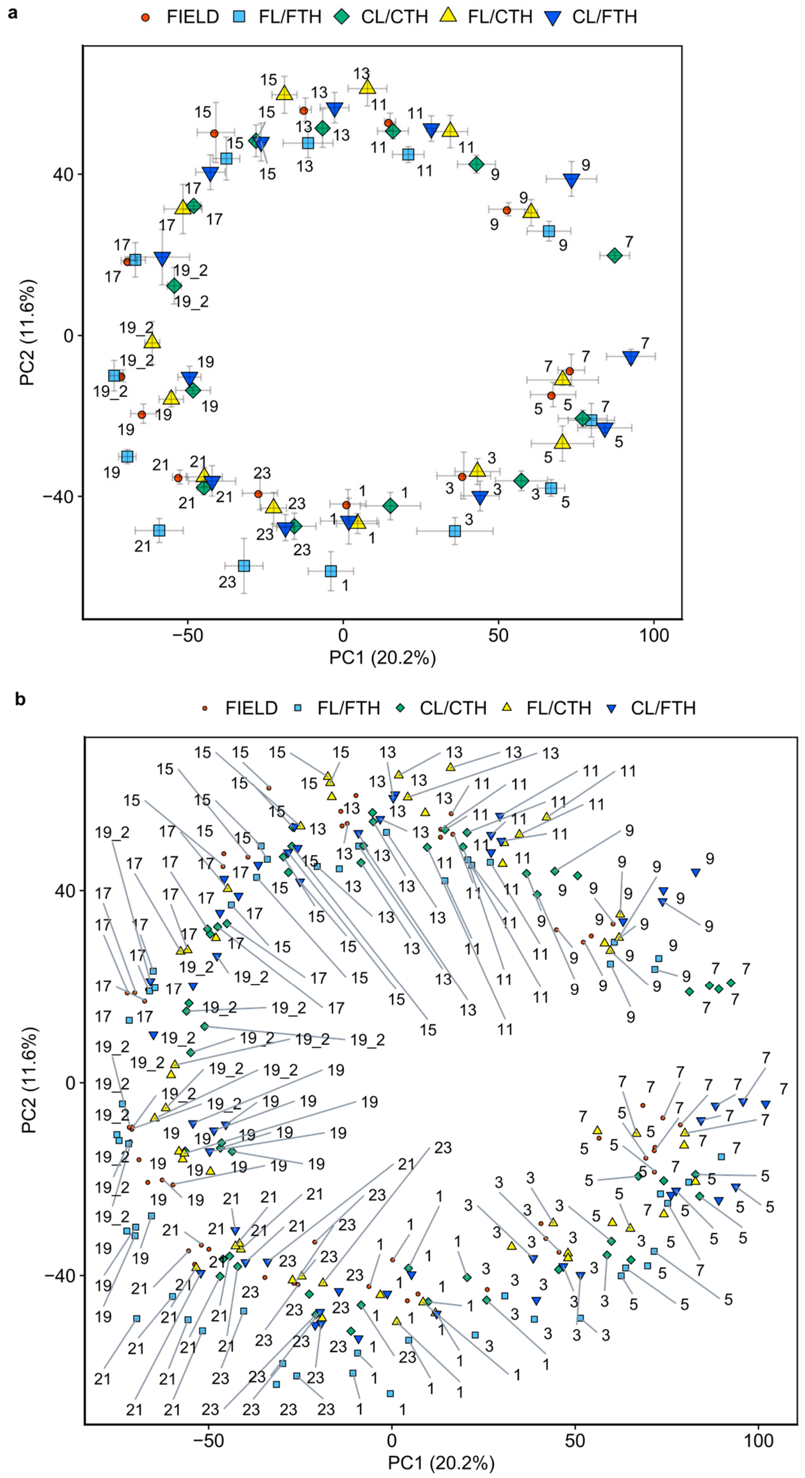


**Supplementary Fig. 6. Principal component analysis (PCA) of transcriptomes in Experiment_1. a**, PCA of transcriptomes at each time point and condition, which corresponds to Fig. 1g. Each point represents the mean value of the four replicates, and error bars indicate the standard errors of PC1 and PC2. **b**, PCA of transcriptomes of each sample at each time point and condition. Four replicates at each time point and condition are shown independently. Numbers indicate sampling times. The percentages of the total variance represented by PC1 and PC2 are shown in parentheses. 19_2 indicates the timepoint 24 h after the start of sampling day at 19:00.

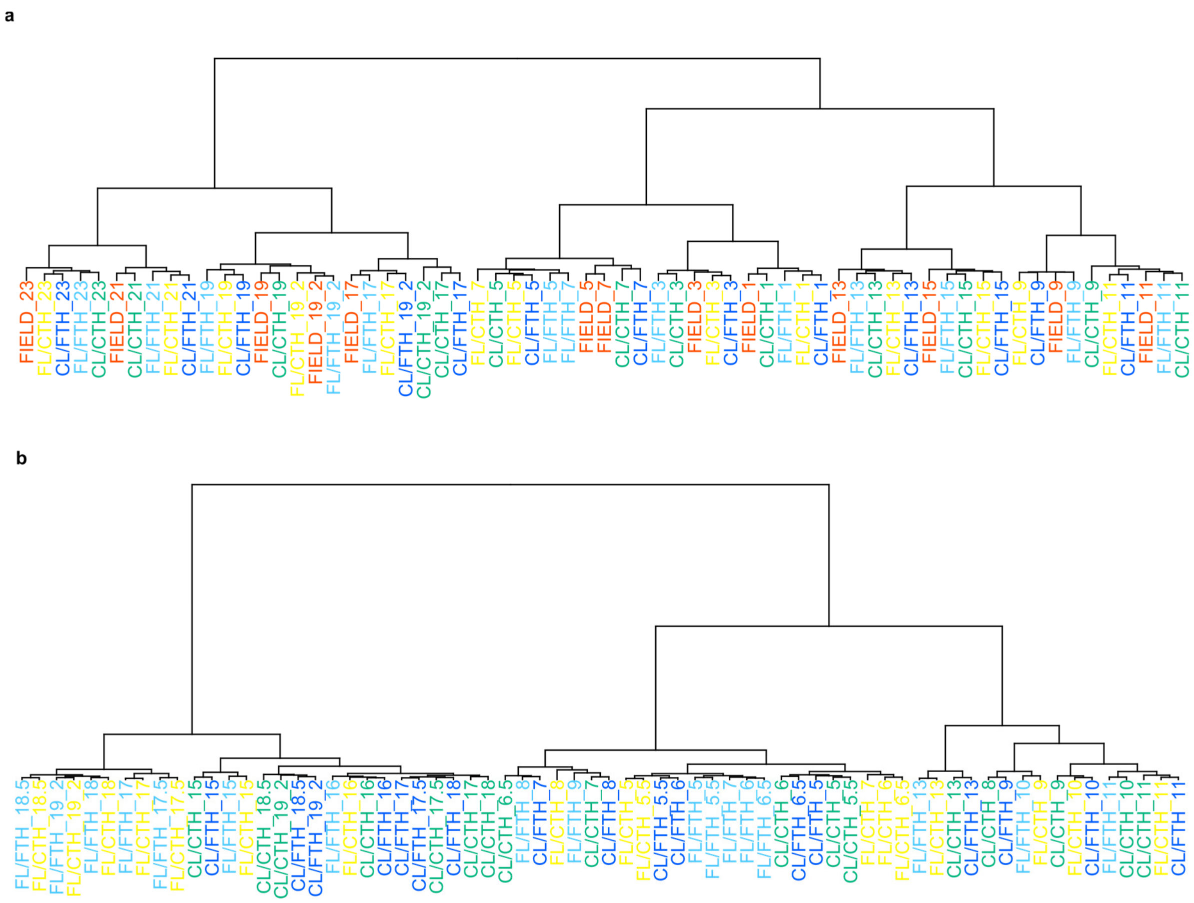


**Supplementary Fig. 7. Hierarchical cluster dendrogram of transcriptomes of each condition and time point.** The dendrograms in Fig. 1i and Fig. 1j correspond to **a** and **b**, respectively. Mean values of (**a**) 4 and (**b**) 3 replicates at each time point and condition were used for the analyses. 19_2 indicates the timepoint 24 h after the start of sampling day at 19:00.

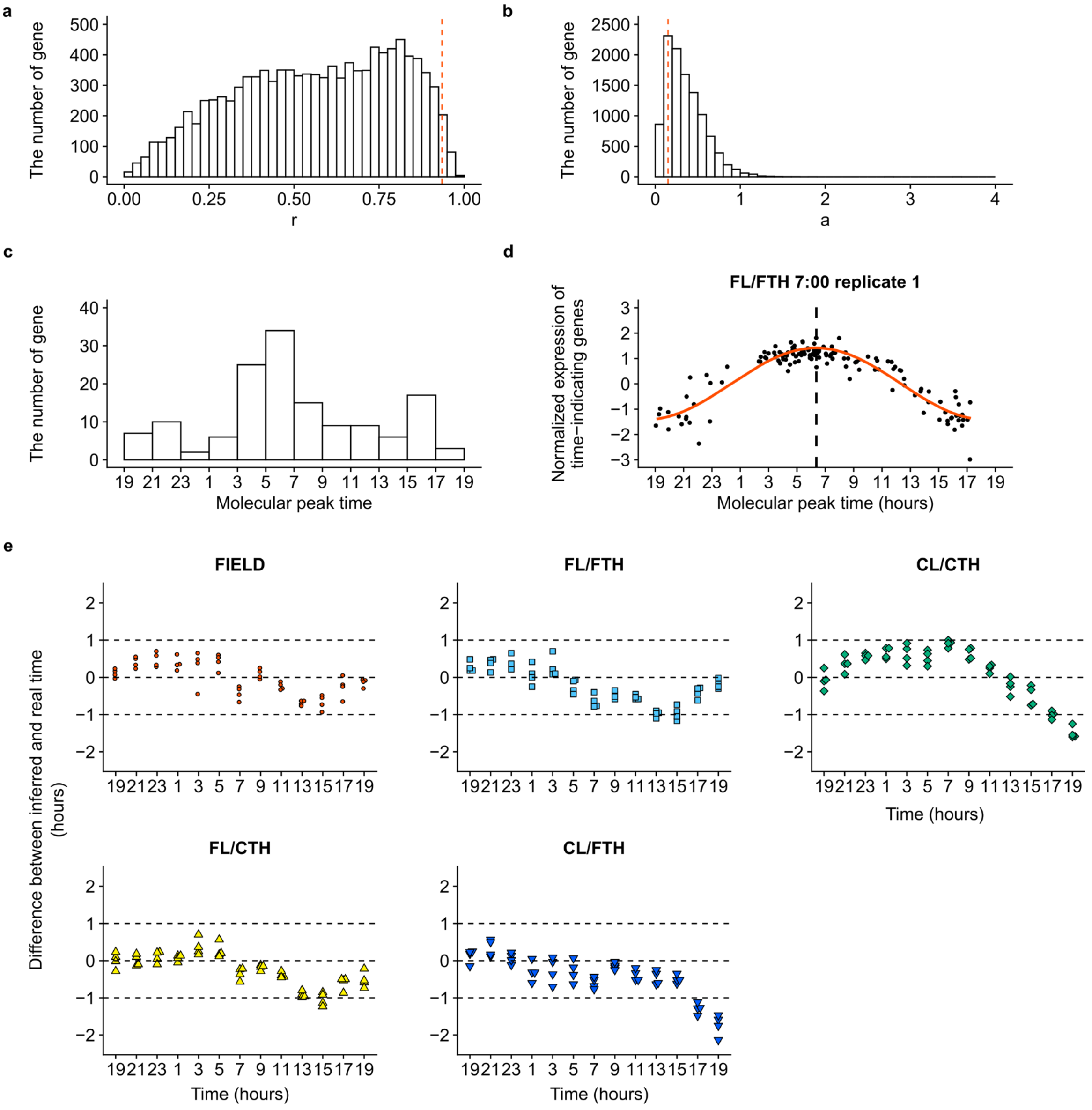


**Supplementary Fig. 8. Inference of internal time using the molecular timetable method.** **a**, Histogram showing the Pearson correlation coefficient (r) between the expression of each gene and its best-fitting cosine curve. The red dashed line indicates the selection threshold (r = 0.935) for time-indicating genes. **b**, Histogram showing the expression amplitude of each gene. The red dashed line indicates the selection threshold (a = 0.15) for time-indicating genes. **c**, Histogram showing the molecular peak times of 143 time-indicating genes. **d**, An example of internal time inference using the molecular timetable method. Normalised expressions of time-indicating genes at the molecular peak time of each gene were fitted to the cosine curve, and the peak time indicates the internal time. **e,** Inference of internal time at each condition in Experiment_1. Inferred time minus real time is shown for each sample (n = 4). Progression of internal time in the evening was slower in CL/CTH and CL/FTH than in FIELD.

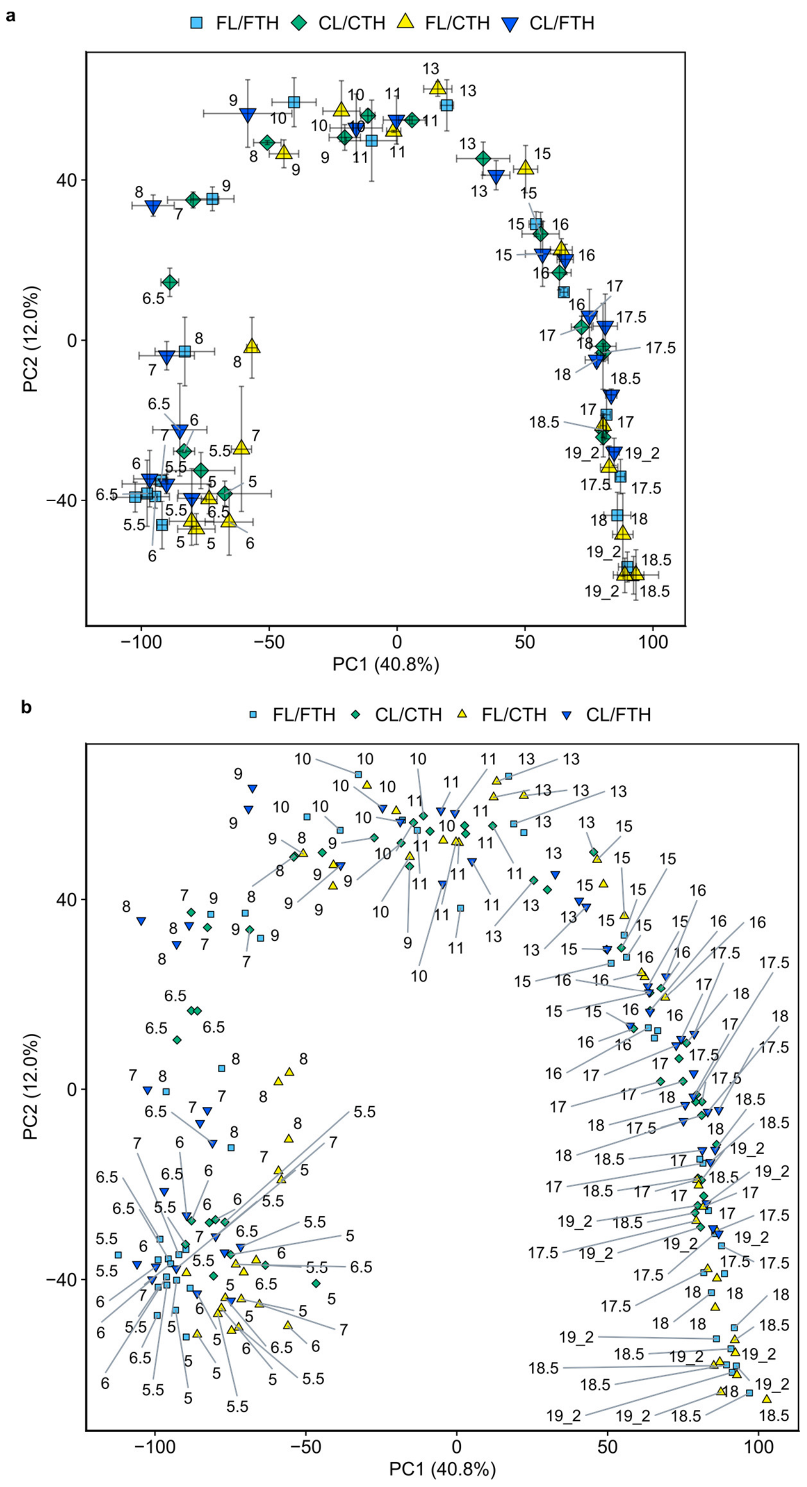


**Supplementary Fig. 9. Principal component analysis (PCA) of transcriptomes in Experiment_2. a**, PCA of transcriptomes at each time point and condition, which correspond to Fig. 1h. Each point represents the mean value of three replicates, and error bars indicate standard errors of PC1 and PC2. **b**, PCA of transcriptomes of each sample at each time point and condition. Four replicates at each time point and condition are shown independently. Numbers indicate sampling times. The percentages of total variance represented by PC1 and PC2 are shown in parentheses. 19_2 indicates the timepoint 24 h after the start of sampling day at 19:00.

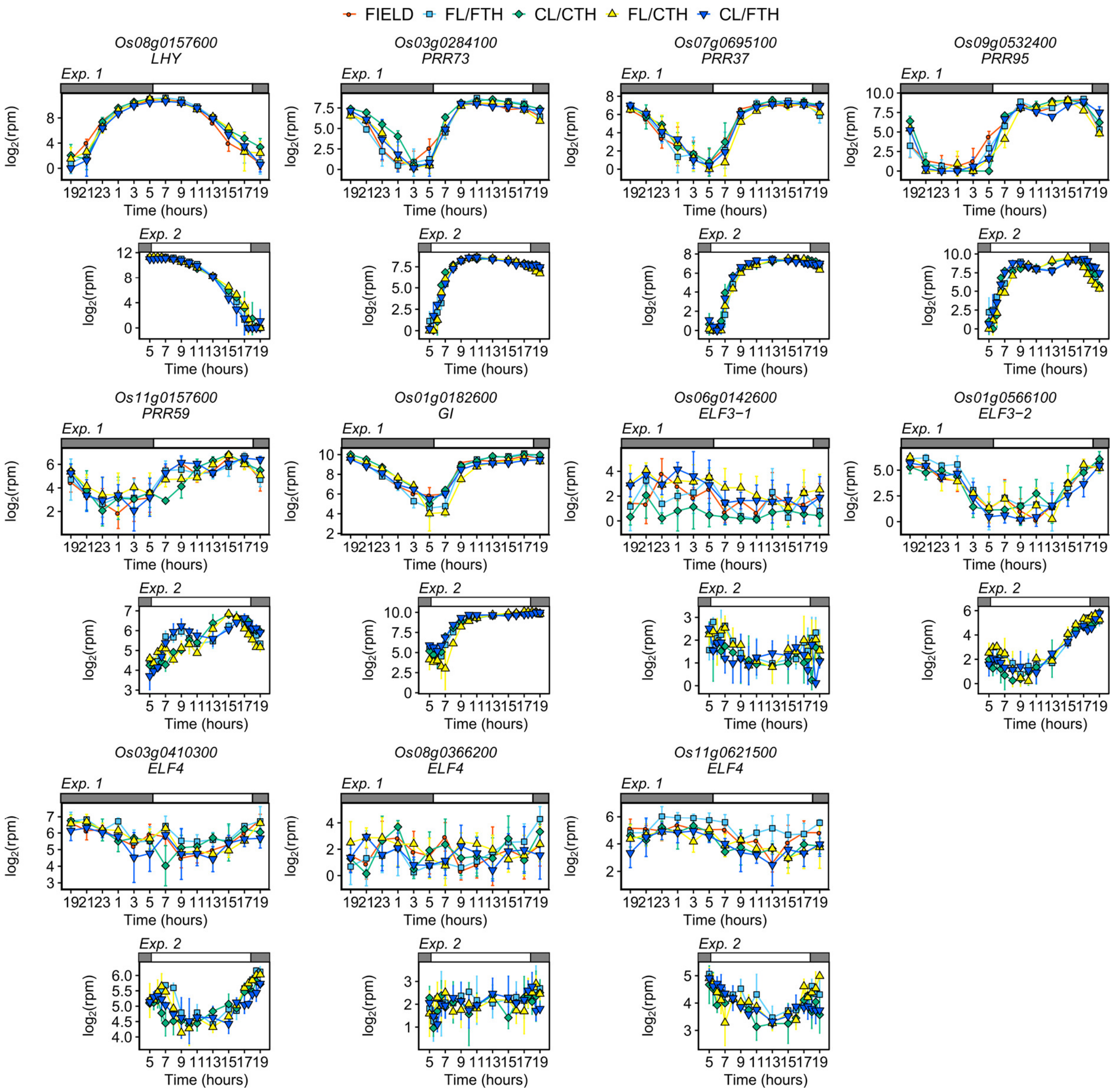


**Supplementary Fig. 10. Expression of genes encoding core circadian clock genes.** Points indicate means and error bars indicate standard deviations (n = 4 and n = 3 in Experiment_1 and Experiment_2, respectively). *LHY*, *LATE ELONGATED HYPOCOTYL*; *PRR73*, *PSEUDO-RESPONSE REGULATOR 73*; *PRR37*, *PSEUDO-RESPONSE REGULATOR 37*; *PRR95*, *PSEUDO-RESPONSE REGULATOR 95*; *PRR59*, *PSEUDO-RESPONSE REGULATOR 59*; *GI*, *GIGANTIA*; *ELF3*, *EARLY-FLOWERING 3*; *ELF4*, *EARLY-FLOWERING 4*.

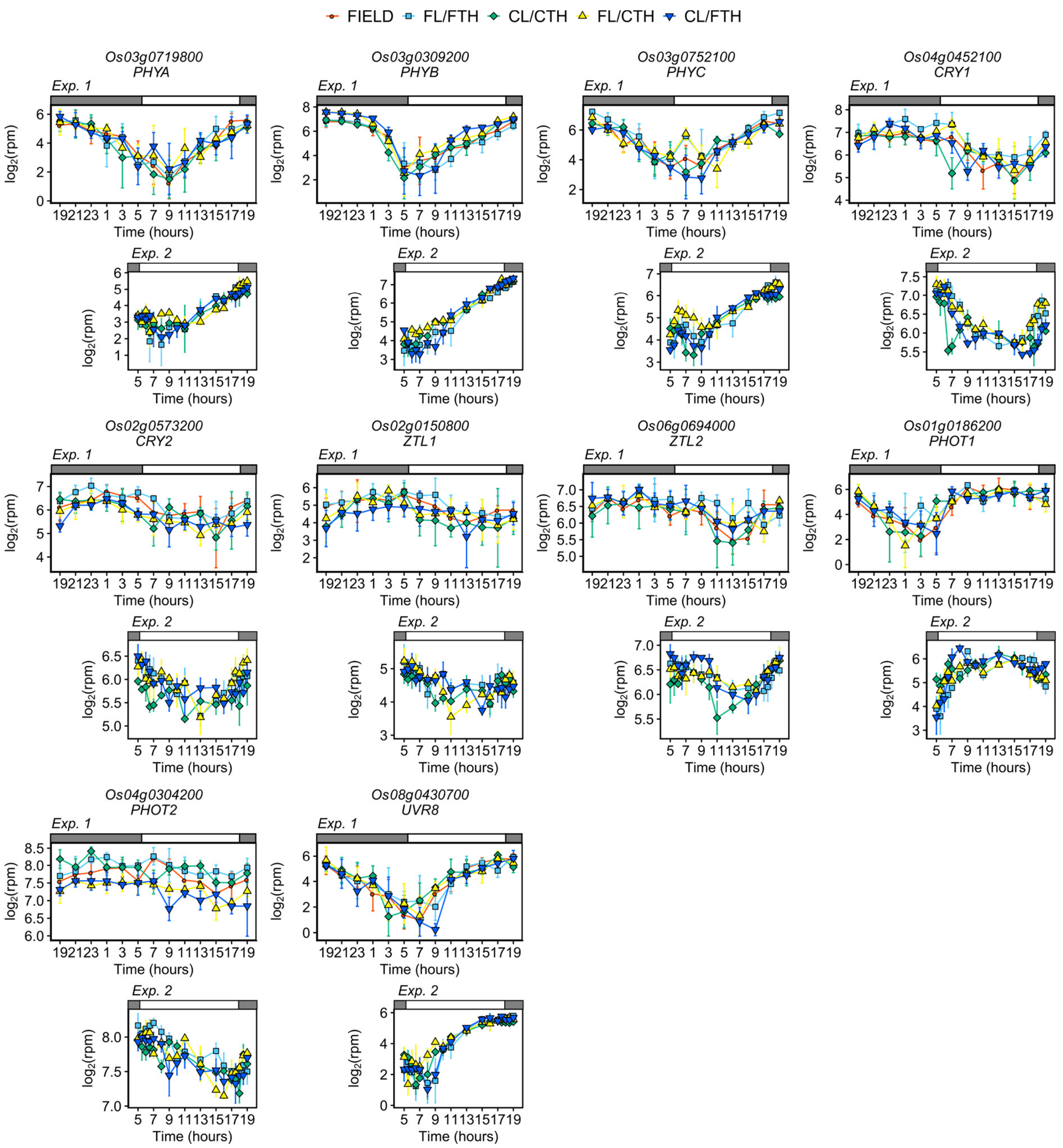


**Supplementary Fig. 11. Expression of genes encoding photoreceptors.** Points indicate means and error bars indicate standard deviations (n = 4 and n = 3 in Experiment_1 and Experiment_2, respectively). CRY, cryptochrome; PHOT, phototropin; PHY, phytochrome; UVR8, UV-B resistance 8; ZTL, *ZEITLUPE*.

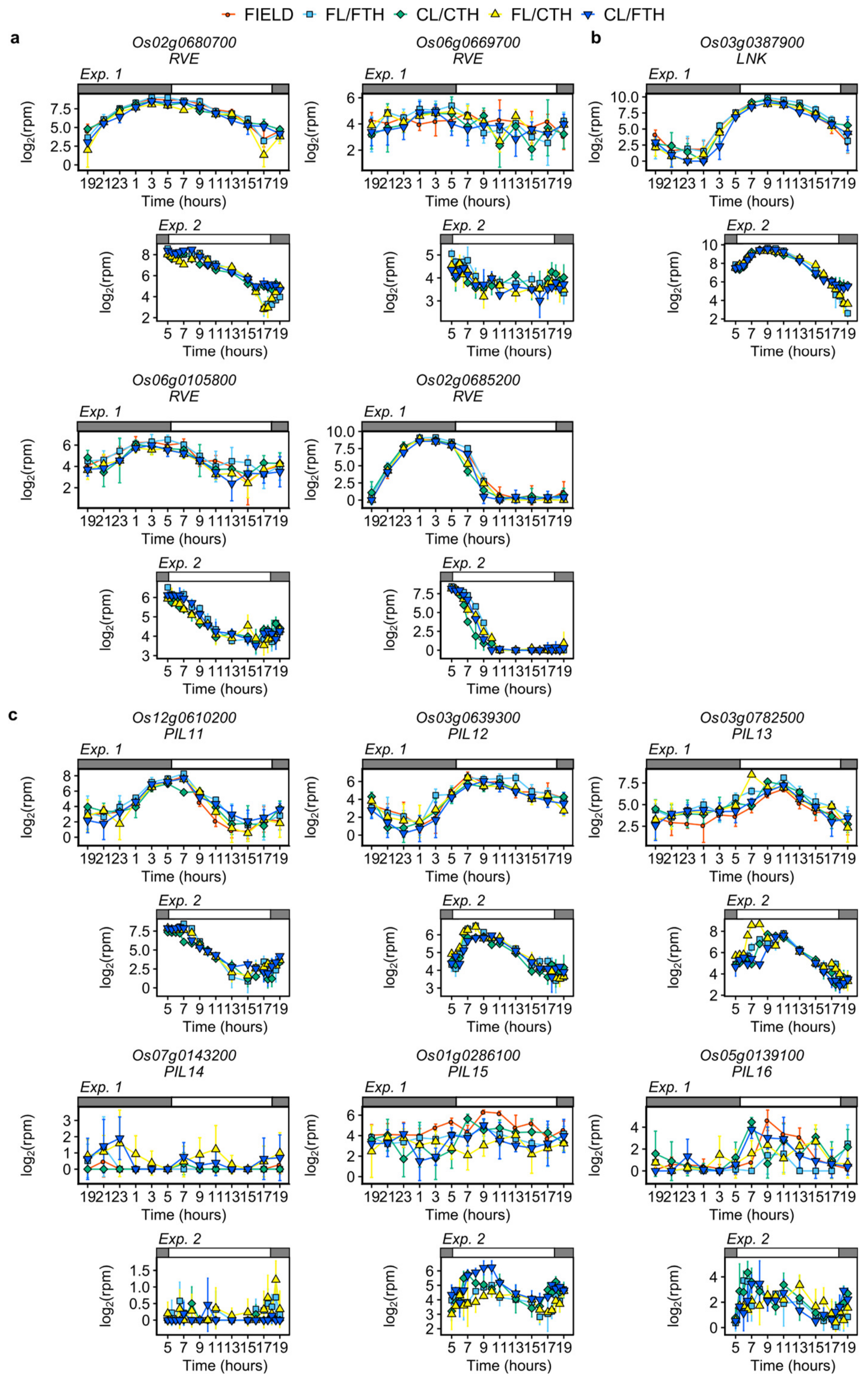


**Supplementary Fig. 12. Expression of genes encoding (a) REVEILLE (RVE), (b) NIGHT LIGHT-INDUCIBLE AND CLOCK-REGULATED protein (LNK), and (c) PHYTOCHROME INTERACTING FACTOR-LIKE protein (PIL).** Points indicate means and error bars indicate standard deviations (n = 4 and n = 3 in Experiment_1 and Experiment_2, respectively).

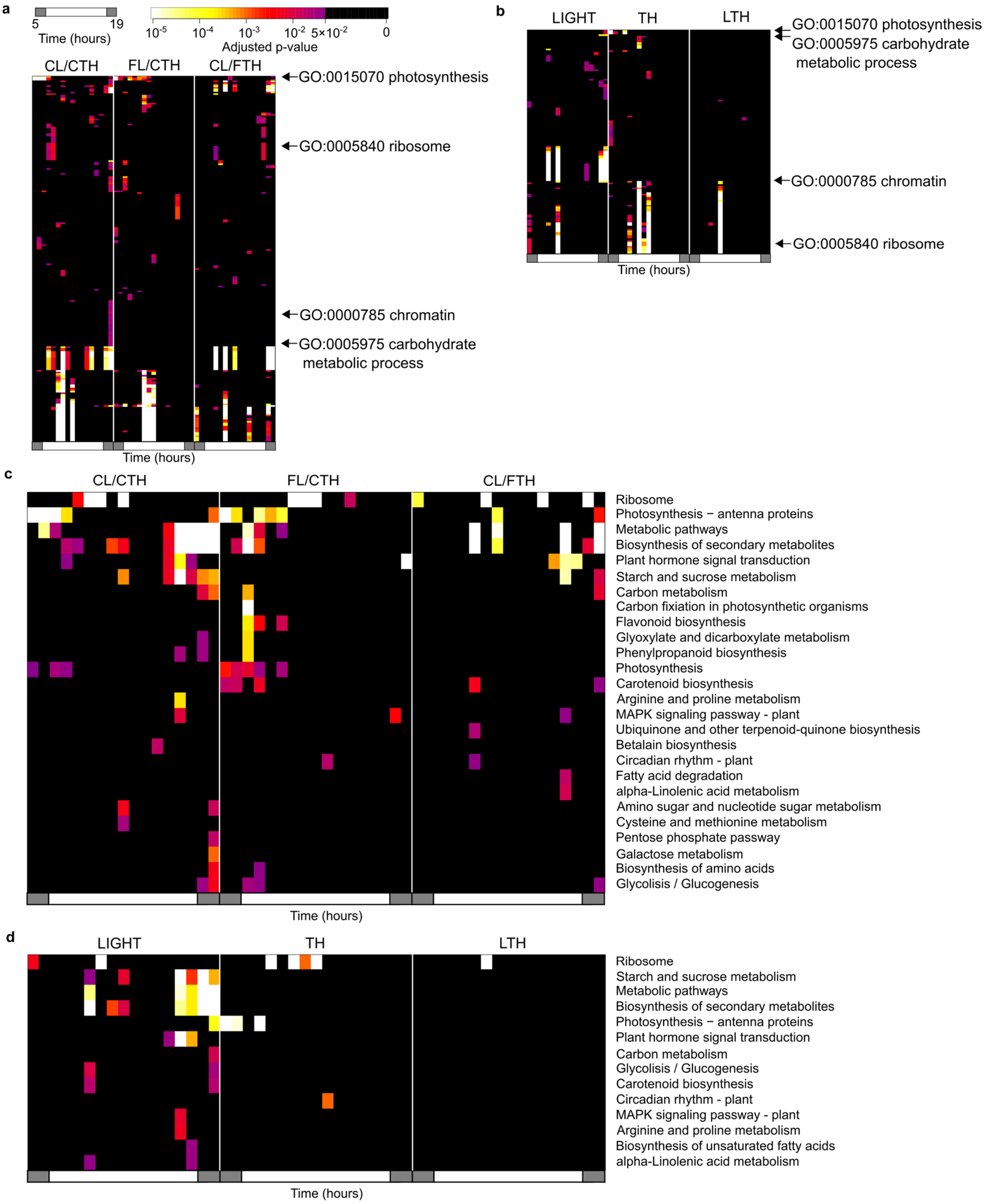


**Supplementary Fig. 13. Significant enrichment of genes with specific annotations in DEGs between FIELD and the other conditions in Experiment_2.** Heatmaps of P-values (Fisher's exact test, two-sided) for significant genes with (**a**, **b**) a particular gene ontology

(GO) and (**c, d**) a particular KEGG pathway (row) in each time and condition (column) in (**a, c**) DEGs between FIELD and the other conditions and (**b, d**) LIGHT, TH, LTH, and UNREP genes. GO and KEGG pathways that have at least one significant (adjusted p-value $< 0.05$) time point and condition are shown.

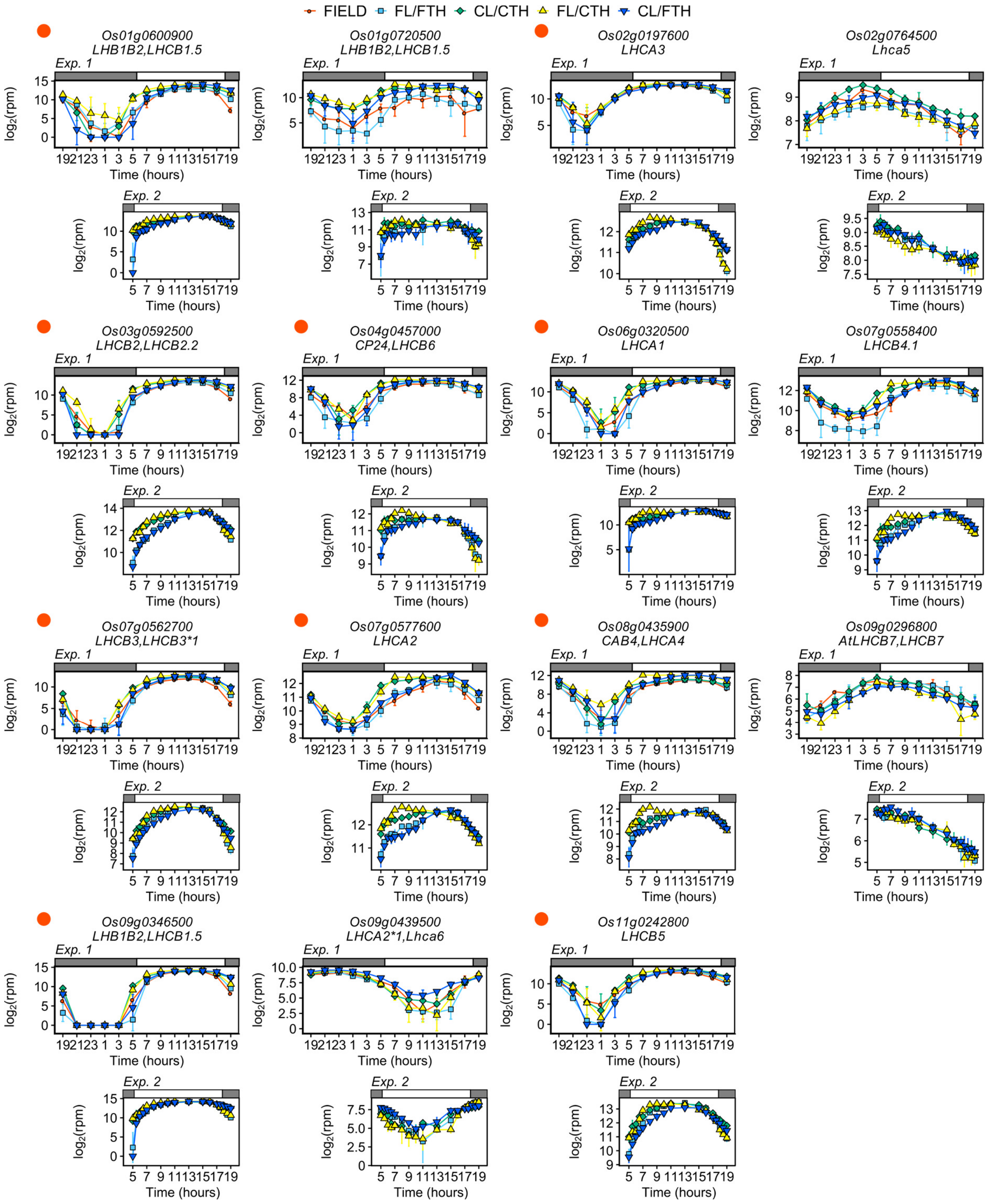


**Supplementary Fig. 14. Expressions of genes related to light harvesting.** Expressions of genes with annotation for photosynthesis-antenna protein (KEGG pathway: dosa00196) or photosynthetic light harvesting (GO:0009765) in Experiment_1 and Experiment_2 are shown.

Red points on the left side of the gene name indicate that the gene was affected by night temperature in Nagano et al.[2] (See Manuscript and Supplemental Table 13). Points indicate means and error bars indicate standard deviations (n = 4 and n =3 in Experiment_1 and Experiment_2, respectively).

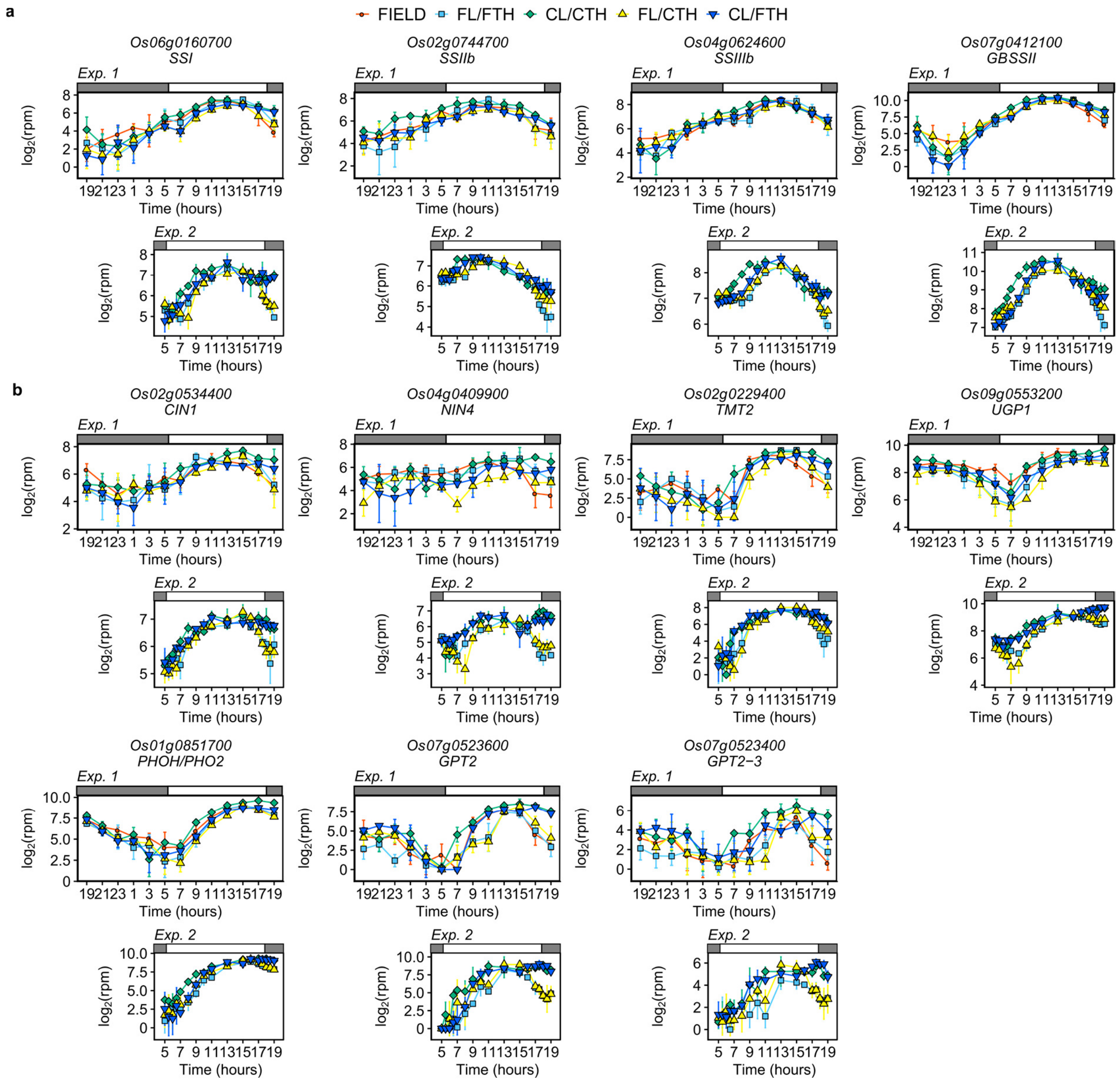


**Supplementary Fig. 15**. **Expression of genes related to (a) starch and (b) sucrose metabolism.** Points indicate means and error bars indicate standard deviations (n = 4 and n = 3 in Experiment_1 and Experiment_2, respectively). *CIN1*, cytosolic invertase 1; *GPT2*, glucose-6-phosphate/phosphate translocator 2; *GPT2-3*, glucose-6-phosphate/phosphate translocator 2-3; *GBSSII*, Granule-bound starch synthase II; *NIN4*, neutral invertase 4; *PHOH/PHO2*, starch phosphorylase 2; *SSI*, starch synthase I; *SSIIb*, starch synthase IIb;

*SSIIIb*, starch synthase *IIIb*; *TMT2*, tonoplast monosaccharide transporter 2; *UGP1*, UDP-glucose pyrophosphorylase 1.

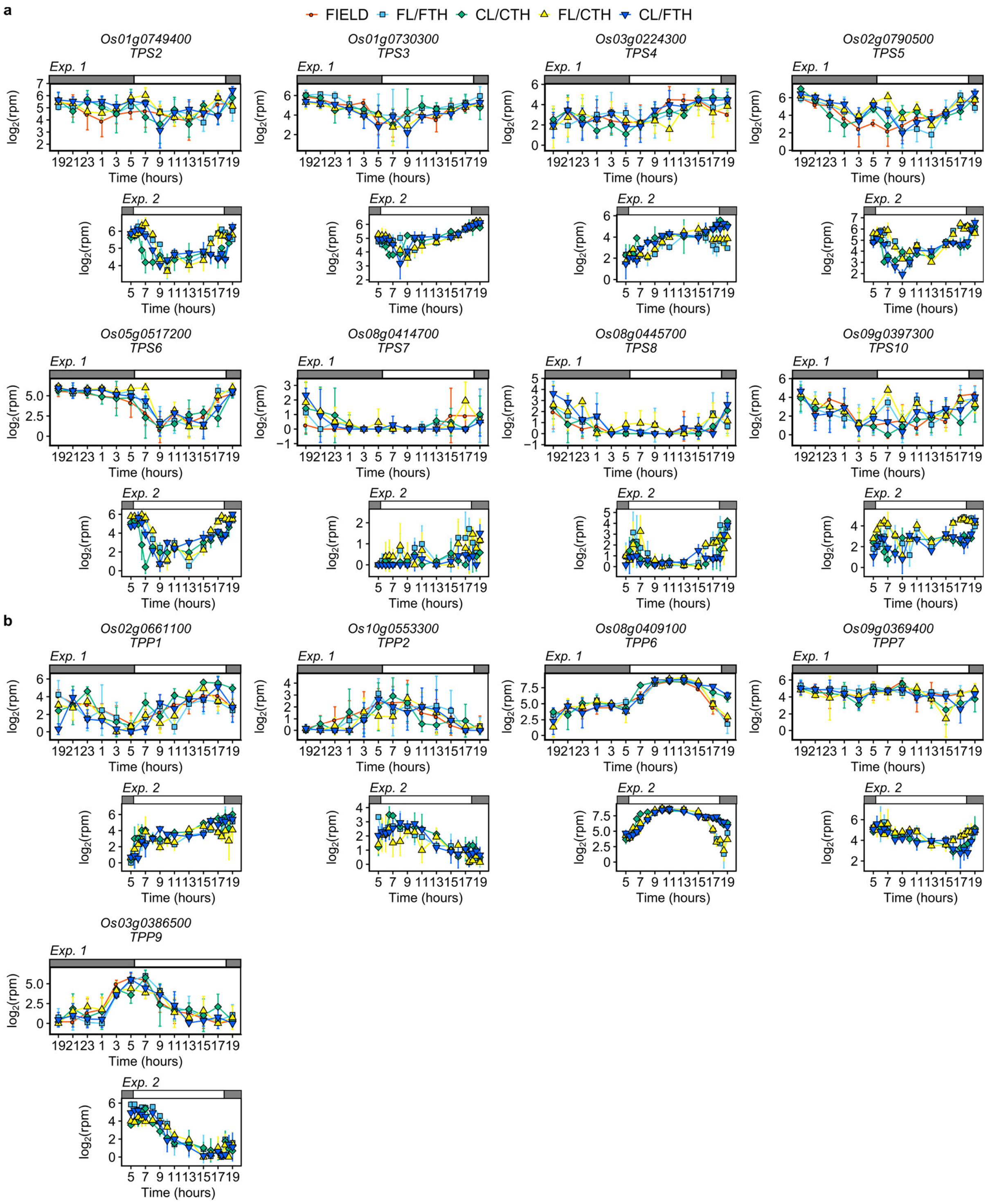


**Supplementary Fig. 16. Expressions of genes encoding (a) trehalose phosphate synthase (TPS) and (b) trehalose phosphate phosphatase (TPP).** Expressions of *TPS1* are shown in Fig. 5e. *TPS9*, *TPS11*, *TPP3*, *TPP4*, *TPP5*, and *TPP8* are not shown because of their low expressions. Points indicate means and error bars indicate standard deviations (n = 4 and n = 3 in Experiment_1 and Experiment_2, respectively).

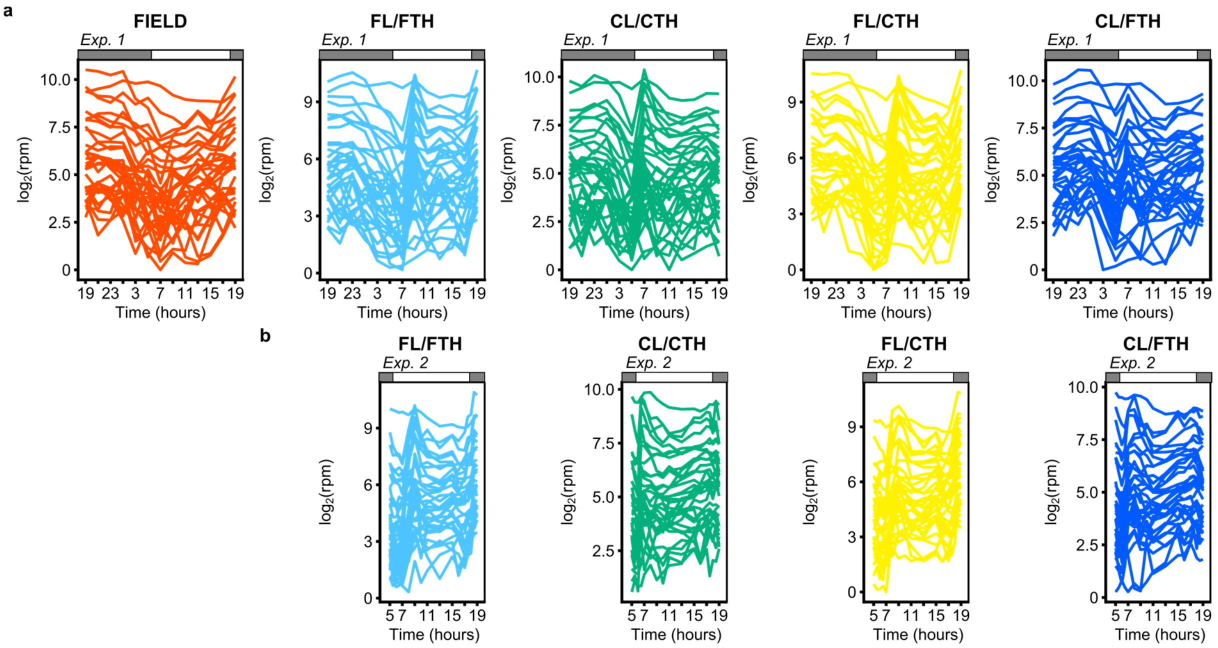


**Supplementary Fig. 17. Different timing of upregulation of chromatin-related genes between conditions.** Expressions of genes with annotations for chromatin (GO:0000785) in **(a)** Experiment_1 and **(b)** Experiment_2. Each line indicates the mean value of the expressions of each gene (n = 4 and n = 3 in Experiment_1 and Experiment_2, respectively). Expressions of genes with annotation for chromatin were induced at 5:00–7:00 in CL/CTH and CL/FTH, and at 7:00–9:00 in FL/FTH and FL/CTH, but the induction in the morning was not obvious in FIELD. The timing of induction was reproduced in Experiment_2. These results suggest that the expression of genes with annotation for chromatin was induced by LED light, but not by sunlight, in the morning, and the timing of induction was affected by diurnal changes in irradiance.

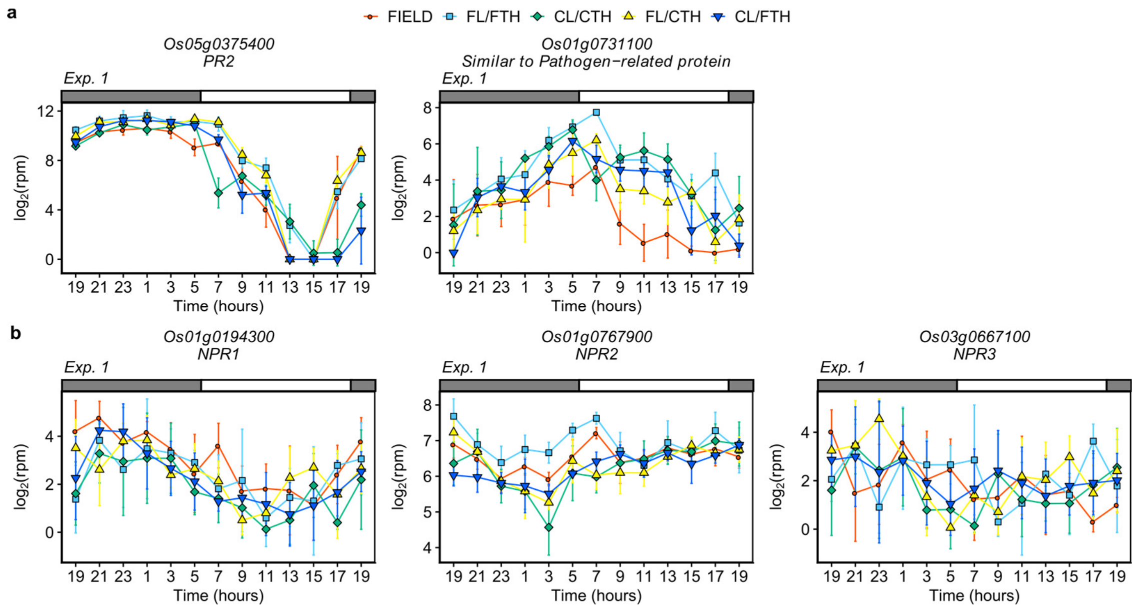


**Supplementary Fig. 18. Expressions of (a) pathogenesis-related (PR) genes and (b) NONEXPRESSOR OF PATHOGENESIS-RELATED (NPR) GENES in Experiment_1.** Points indicate means and error bars indicate standard deviations (n = 4).

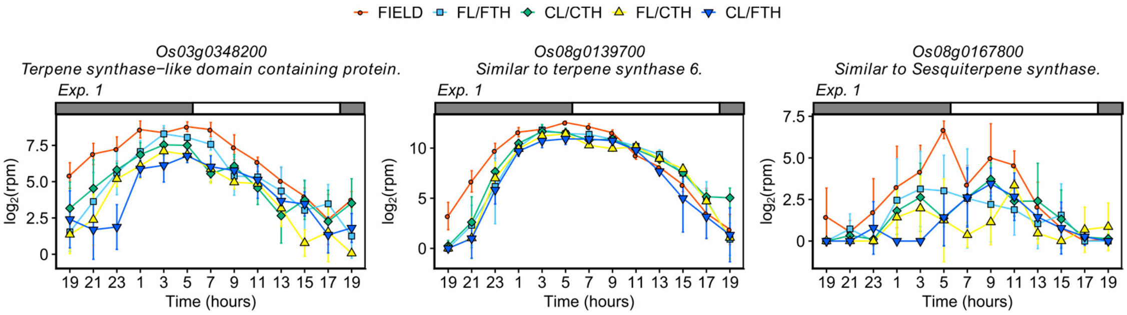


**Supplementary Fig. 19. Expressions of genes related to terpene synthase in Experiment_1.** Points indicate means and error bars indicate standard deviations (n = 4).

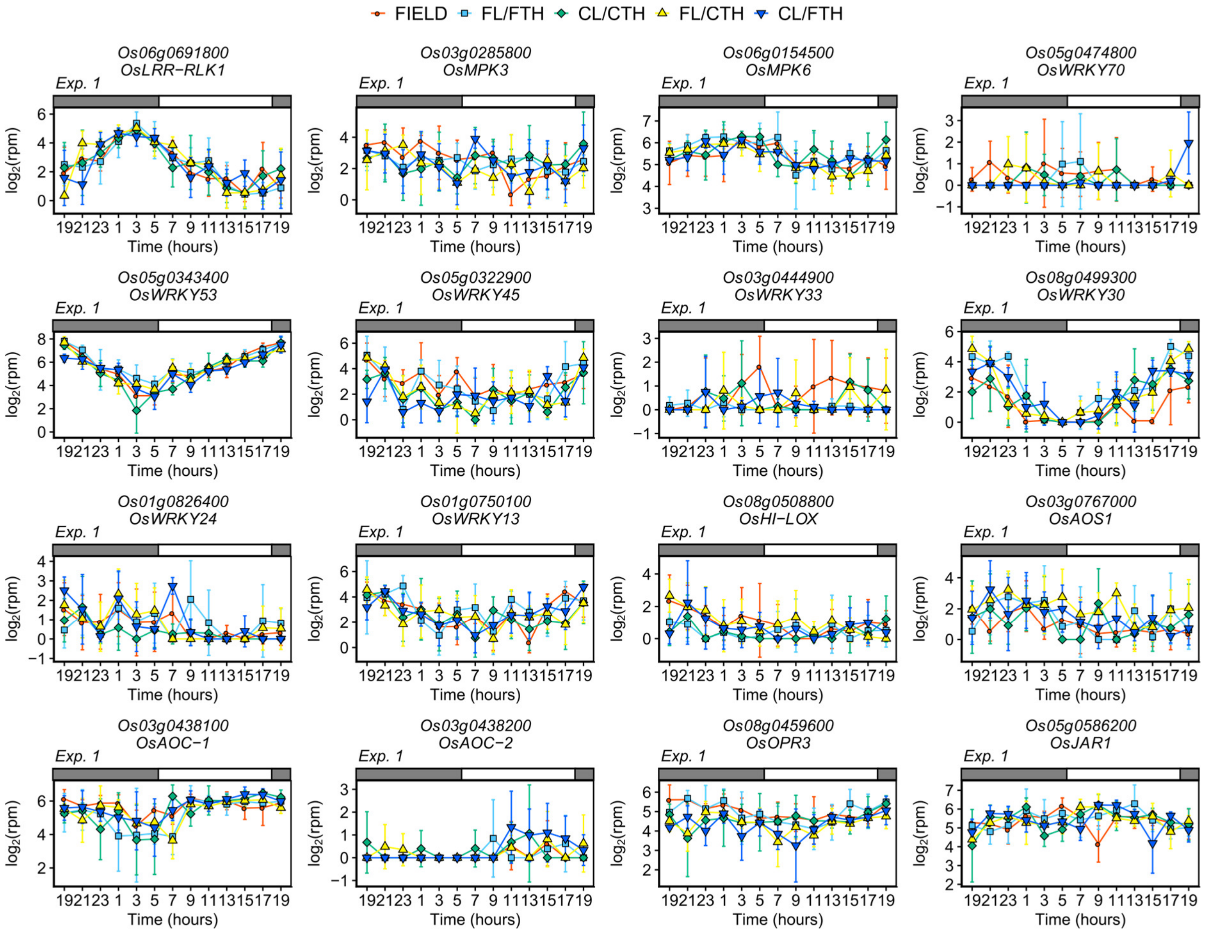


**Supplementary Fig. 20. Expressions of genes involved in early defence signalling to herbivore attack listed in Ye et al.[39] in Experiment_1.** Points indicate means and error bars indicate standard deviations (n = 4). Early defence signalling genes include *Oryza sativa* leucine-rich repeat receptor-like kinase 1 (*OsLRR-RLK1*), mitogen-activated protein kinase (*OsMPK3* and *OsMPK6*), WRKY transcription factors (*OsWRKY13*, *OsWRKY24*, *OsWRKY30*, *OsWRKY33*, *OsWRKY45*, *OsWRKY53*, and *OsWRKY70*) and jasmonate synthesis genes (*OsHI-LOX*, *OsAOS1*, *OsAOC*, *OsOPR3*, and *OsJAR1*).

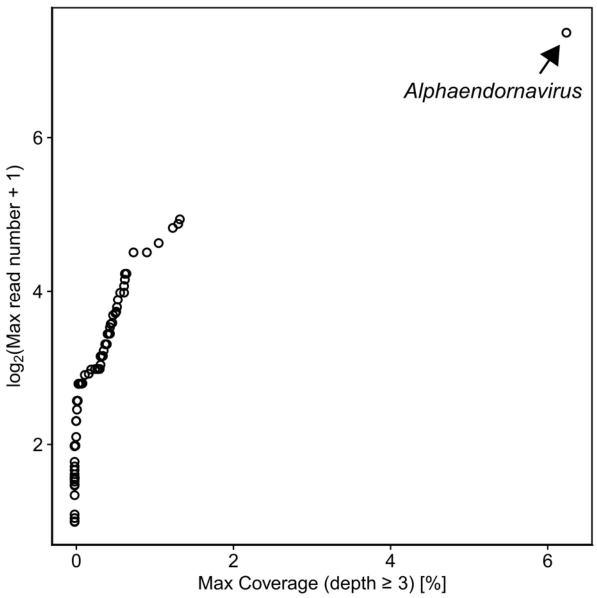


**Supplementary Fig. 21. Detection of virus infection from RNA-Seq data.** Scatter plot showing the max read number and max coverage of 115 viruses with maximum reads per sample >1. *Alphaendornavirus*, which shows no clear symptoms in rice plants[41], had the highest reads (maximum 166 reads per sample) and coverage of the reads to the reference (6.3% with depth ≥ 3). Correlations between the read number of the *Alphaendornavirus* and the expressions of PR genes are shown in Fig. 6f. The number of reads and the coverage of the *Alphaendornavirus* were low in comparison with infected samples discussed in Kamitani et al.[40] (more than 10% in a sample with the lowest coverage). Therefore, this result suggests that even if *Alphaendornavirus* existed in the rice leaves, the copy number of the virus was limited.

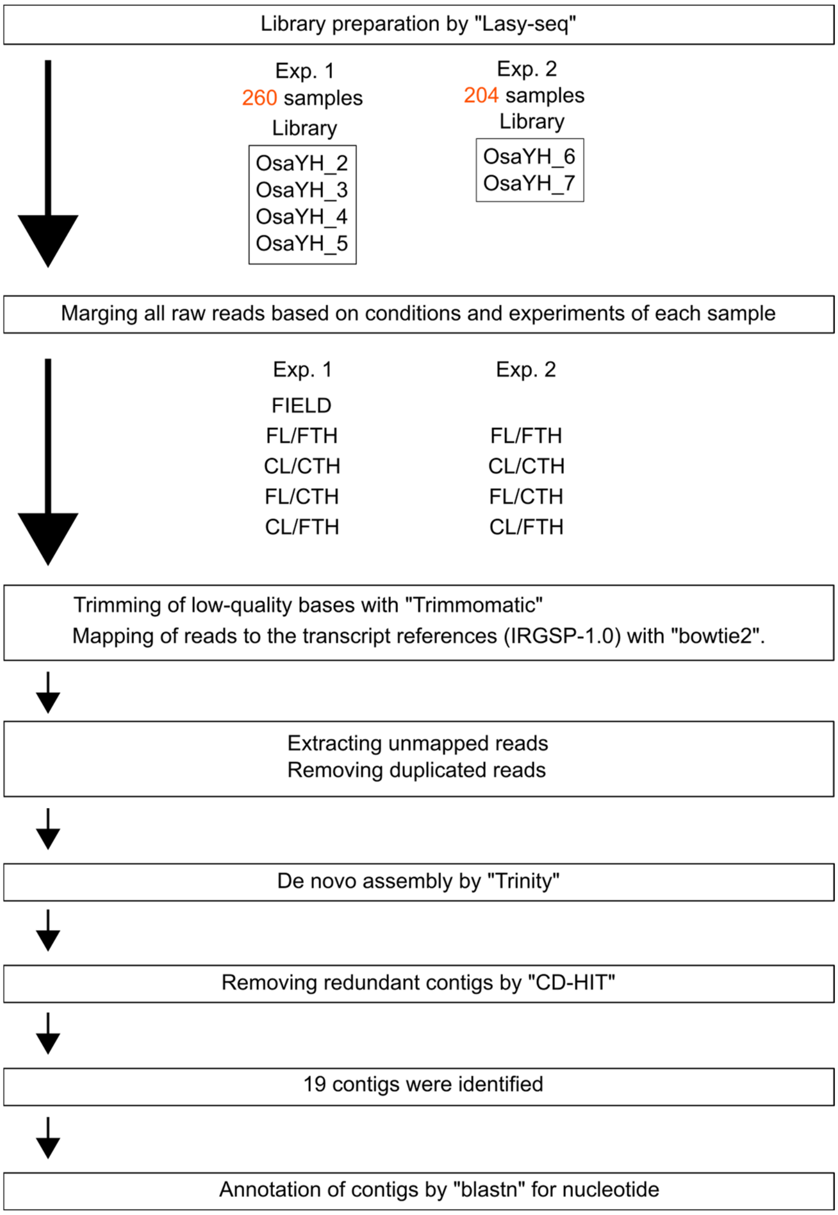


**Supplementary Fig. 22. Scheme for searching for genes that were found by de novo transcriptome assembly from unmapped reads to the rice reference transcriptomes.**

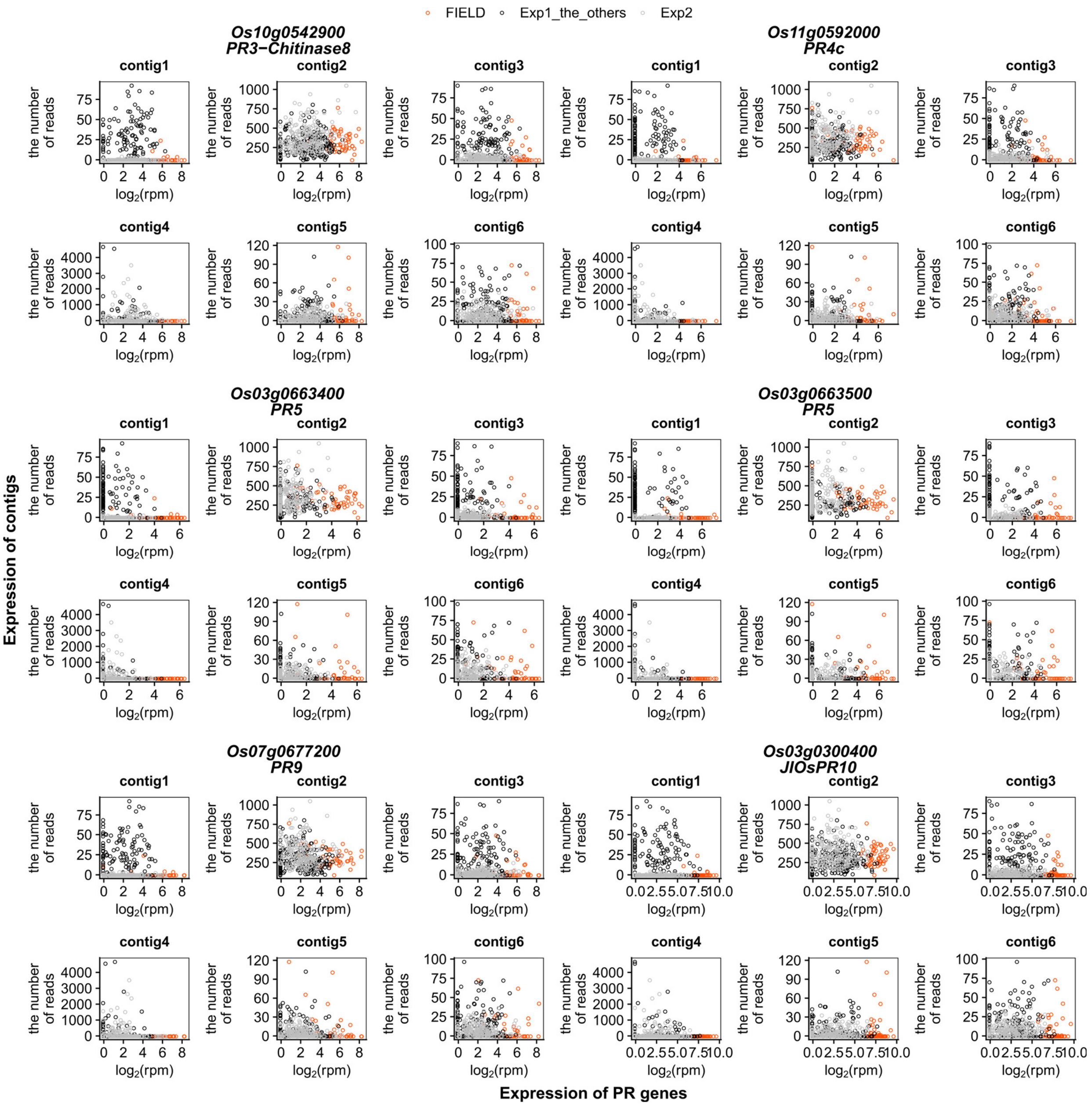


**Supplementary Fig. 23. Scatter plot showing the relationship between PR genes and contigs found by de novo transcriptome assembly from unmapped reads to the rice reference transcriptomes.** We selected 6 contigs from 19 contigs after excluding those with annotations for *Alphaendornaviruses*, synthetic constructs, and expression vectors. contig1, exp1_CL/FTH.TRINITY_DN138_c0_g1_i1; contig2, exp1_FL/CTH.TRINITY_DN11_c0_g1_i1; contig3, exp1_FL/CTH.TRINITY_DN249_c0_g1_i2; contig4, exp2_CL/CTH.TRINITY_DN586_c0_g1_i1; contig5, exp2_CL/FTH.TRINITY_DN544_c0_g3_i1; contig6, exp2_FL/CTH.TRINITY_DN119_c0_g1_i1. Details of each gene are shown in Supplementary Table 17.

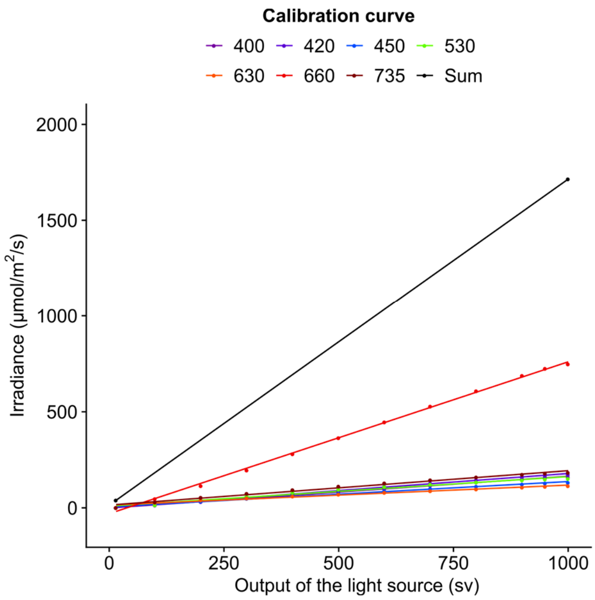


**Supplementary Fig. 24. Calibration curve of the output of the light source of SmartGC versus irradiance.** Calibration curve for seven types of LED light and the sum of all LEDs are shown. The unit of irradiance is photon flux density (PFD) defined over 380–780 nm.
